# Shape matters: the relationship between cell geometry and diversity in phytoplankton

**DOI:** 10.1101/2020.02.06.937219

**Authors:** Alexey Ryabov, Onur Kerimoglu, Elena Litchman, Irina Olenina, Leonilde Roselli, Alberto Basset, Elena Stanca, Bernd Blasius

## Abstract

Size and shape profoundly influence an organism’s ecophysiological performance and evolutionary fitness, suggesting a link between morphology and diversity. However, not much is known about how body shape is related to taxonomic richness, especially in microbes. Here we analyse global datasets of unicellular marine phytoplankton, a major group of primary producers with an exceptional diversity of cell sizes and shapes and, additionally, heterotrophic protists. Using two measures of cell shape elongation, we quantify taxonomic diversity as a function of cell size and shape. We find that cells of intermediate volume have the greatest shape variation, from oblate to extremely elongated forms, while small and large cells are mostly compact (e.g., spherical or cubic). Taxonomic diversity is strongly related to cell elongation and cell volume, together explaining up to 92% of total variance. Taxonomic diversity decays exponentially with cell elongation and displays a log-normal dependence on cell volume, peaking for intermediate-volume cells with compact shapes. These previously unreported broad patterns in phytoplankton diversity reveal selective pressures and ecophysiological constraints on the geometry of phytoplankton cells which may improve our understanding of marine ecology and the evolutionary rules of life.

## Introduction

High diversity of organismal body shapes evolved as a result of natural selection of morphological traits in response to variable environmental conditions, interactions with other species, and availability of ecological niches. There is a large body of literature on the effects of external factors on body shape and the effects of body shape on species fitness. However, these studies mostly focus on complex animals, while body shape patterns and their effects on fitness in the most diverse ecological groups – unicellular organisms – are much less studied. Furthermore, the analyses have been typically focused on the effects of certain environmental conditions on cell shape distribution, and little is known about the overall diversity of shape classes, their distribution and the effects of body shape on taxonomic diversity, and the ultimate evolutionary success of organisms of a given shape. Here we analyse the size and shape distributions of unicellular marine photosynthetic microbes – major aquatic primary producers forming the base of most marine food webs, with the addition of some heterotrophic forms (dinoflagellates). We discuss various approaches to characterize cell shape variation, analyse the diversity of shape classes and investigate how taxonomic diversity varies across cell volume and cell shape classes.

### Body shape adapts to environment

Adaptation to the physical and ecological environment is a key evolutionary process. As Darwin pointed out more than 150 year ago, species can lose or gain morphological traits as a result of natural selection (Darwin 1859). Body shape affects metabolic rates (Hirst *et al*. 2014), and can rapidly adapt to the environment (Husemann *et al*. 2017). For instance, contrasting physical environments lead to differences in limb length between aquatic and terrestrial salamanders (Edgington & Taylor 2019), temperature changes caused either by latitudinal gradient or global warming affect shapes of lizards (Forsman & Shine 1995)various endotherms (Porter & Kearney 2009) and birds’ wing size (Weeks *et al*. 2020), and fishing creates anthropogenic selection pressure on body shape distribution of diverse fishes (Alós *et al*. 2014).

Thus, if a certain body shape matches some environmental conditions better than others, then organisms with that body shape should have higher fitness and, consequently, diversity. However, the effect of body shape on diversity is largely unexplored. Most studies have focused on the effects of body size, and showed that body size is typically a strong predictor of biodiversity: a unimodal relationship between body size and species richness was demonstrated across various taxa (May 1986), including insects (Siemann *et al*. 1996), reptiles (Feldman *et al*. 2016), mammals (Martin 2017), and fishes (Albert & Johnson 2012). In phytoplankton, the patterns are less clear: some found that local species richness can vary as a hump-shaped function of volume (Cermeño & Figueiras 2008), whereas other studies indicate that it decreases as a power function of volume (Ignatiades 2017). Moreover, the effects of body shape on phytoplankton taxonomic diversity remain unexplored.

### Unexplored diversity and role of phytoplankton cell shapes

Phytoplankton is vital to the functioning of marine ecosystems. It is exposed to a continuously changing environment. A large array of continually varying environmental factors drives phytoplankton growth, with the vertical and horizontal resource gradients, as well as the dynamics of predator and nutrient distributions giving rise to a myriad of ecological strategies and rich taxonomic diversity (Hutchinson, 1961). Water is in constant movement due to winds, currents, tides, and temperature changes, and the phytoplankton species need to be adapted to a wide range of hydrodynamic conditions. However, for microorganisms this adaptation is different from macroorganisms, because for microbes the ratio between inertial and viscous forces, the so-called Reynolds number, is much smaller than one. This means that, while the main physical forces acting on macroorganisms are inertial and proportional to body weight, the main forces acting on microorganisms are viscous and proportional to the cell surface area (Naselli-Flores *et al*. 2020).

Cell shape and size are the main factors determining total volume and surface area, with cell volumes spanning many orders of magnitude and dozens of different shape types, from simple spherical to extremely complex and elongated cells (Fig. 1). The variation in shape results in a non-linear relationship between cell linear dimensions and volume (Mittler *et al*. 2019). The shape of planktonic cells varies in two main aspects: the *geometric* shape of a cell (spherical, elliptic, conical, etc.) and the shape *elongation* or *flattening*. Together with cell size, these factors are the most easily measured cell traits for automated monitoring systems (Pomati *et al*. 2011) and can be used as key morphological traits (Naselli-Flores & Barone 2011; Stanca *et al*. 2013).

**Fig. 1.**
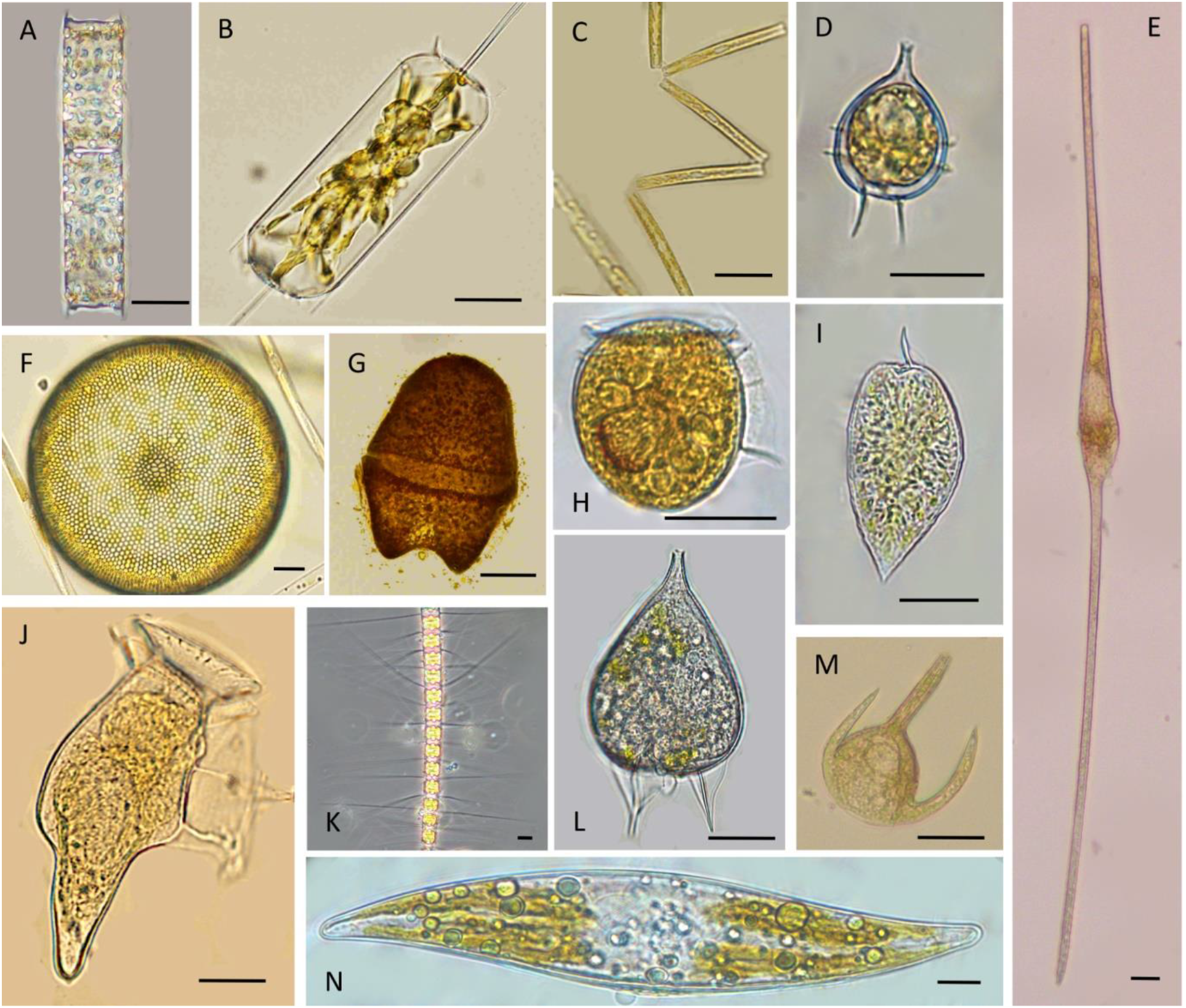
Examples of phytoplankton cells, their relative shape and elongation class. (A) *Cerataulina pelagica* (Cylinder, prolate); (B) *Ditylum brightwellii* (prism on triangular base, prolate); (C) *Thalassionema nitzschioides* (parallelepiped, prolate); (D) *Protoperidinium* sp. (cone + half sphere, compact); (E) *Tripos fusus* (double cone, prolate); (F) *Coscinidiscus* sp. (cylinder, oblate); (G) *Akashiwo sanguinea* (ellipsoid, oblate); (H) *Phalacroma* sp. (ellipsoid, oblate); (I) *Prorocentrum micans* (cone + half sphere, prolate); (J) *Dinophysis caudata* (ellipsoid + cone, prolate); (K) *Chaetoceros didymus* (prism on elliptic base, oblate); (L) *Podolampas bipes* (cone, compact); (M) *Tripos* sp. (ellipsoid + 2 cones + cylinder, prolate); (N) *Pleurosigma* sp. (prism on parallelogram base, prolate). Scale bar = 20 µm.

It is known that environmental conditions, such as nutrients, light, temperature and grazers, affect the shape and size distributions of phytoplankton (Naselli-Flores et al., 2007; Stanca et al., 2013; Zohary et al., 2010), confirming that both size and shape are crucial determinants of fitness. Environmental factors can select specific phytoplankton morphological groups (Kruk & Segura 2012). The seasonal patterns of morpho-functional groups are often similar across different lakes (Naselli-Flores & Barone 2007) and different years (Weithoff & Gaedke 2017), even though the precise species composition might differ (Salmaso & Padisák 2007; Hillebrand *et al*. 2018). Consequently, the composition of morphological traits might have a greater consistency than species composition.

Cell shape affects many aspects of phytoplankton survival, such as grazing by zooplankton (Sunda & Hardison 2010; Pančić & Kiørboe 2018), diffusion and sinking (Padisák *et al*. 2003; Durante *et al*. 2019), maximal growth rates (Wirtz 2011), nutrient uptake (Grover 1989; Karp-Boss & Boss 2016) and harvesting of light (Naselli-Flores & Barone 2007, 2011). Because the surface to volume ratio (*S*/*V*) decreases with cell volume, it has been often assumed that for cells of large volumes, natural selection should favor elongated or flattened shapes with increased surface area (Lewis 1976; Niklas 2000). This can increase the number of nutrient acquisition sites on the surface, maximizing nutrient uptake (Karp-Boss & Boss 2016) or improving chloroplast packing, minimizing shading and increasing light harvesting (Naselli-Flores & Barone 2007, 2011). However, direct experimental support of this is scarce. By contrast, it is known that for many animals, resource uptake in 3D environments scales linearly with body size (Pawar *et al*. 2012). For phytoplankton, nutrient uptake rates and quotas typically scale proportionally to volume or carbon content (Edwards *et al*. 2012; Dao 2013; Marañón 2015), with large cells often having compact shapes.

### Exploring shape-diversity relationships

The large variation in phytoplankton cell volumes and shapes, and their dependence on environmental conditions, present a unique opportunity to investigate evolutionary constraints on morphological traits and their connection to taxonomic diversity. Using the most comprehensive dataset of phytoplankton cell sizes and shapes, we address several novel questions. We determine if there are broad patterns in cell volume and shape variation of marine unicellular organisms across main phyla of phytoplankton and heterotrophic dinoflagellates (together called below, for brevity, phytoplankton). We ask whether some shapes and combinations of cell size and shape are more common than others and whether the patterns are similar across different phyla. We explore whether certain shapes lead to a greater diversification, resulting in higher taxonomic richness and whether there is a relationship between cell shape and taxonomic richness, and if it can be predicted from fundamental constraints on cell dimensions.

## Methods

### Data sources

We compiled a comprehensive data sets of phytoplankton and other marine protists in terms of sizes, shapes and taxonomic diversity from seven globally distributed marine areas: Baltic Sea, North Atlantic (Scotland), Mediterranean Sea (Greece and Turkey), Indo-Pacific (the Maldives), South-western Pacific (Australia), Southern Atlantic (Brazil). The data comprise 5,743 cells of unicellular phytoplankton from 402 genera belonging to 16 phyla identified according to www.algaebase.org (Guiry & Guiry 2018).

The data sources include two datasets. The first dataset represents the results of monitoring in several stations in Baltic Sea over the past 25 years (with interval 1-2 months from May to November) and contains information on phytoplankton species and heterotrophic dinoflagellates covering a total of 308 genera. The second dataset includes a biogeographical snapshot survey of phytoplankton assemblages obtained by Ecology Unit of Salento University performed during summer in 2011 and 2012 in six coastal areas with different biogeographical conditions (ecoregions) around the globe (Roselli *et al*. 2017). This survey included 3 concurrent data replicas from each of 116 local sites. This data covers a total of 193 genera sampled from 23 ecosystems of different typology (coastal lagoons, estuaries, coral reefs, mangroves and inlets or silled basins). The data used in this study are available online (ICES CEIM; LifeWatch ERIC), see also Data availability for the data included in manuscript submission.

The datasets were obtained using different techniques and over different time intervals. The regular (with 1-2 month intervals) monitoring of plankton in Baltic sea was performed over the past 25 years, at the same stations and includes data for cells less than 1 *μm*in length, while for the second dataset phytoplankton was sampled only once per location, but in various regions of the world ocean and with mesh size of 6 *μm*(See Sampling methods below). The first dataset includes a wider range of cell volumes from 0.065 *μ*m^3^ to 5 · 10^8^ *μ*m^3^ and more species, and the second dataset represents only a part of this size distribution in the range of volumes from 5.9 *μ*m^3^ to 3.9 · 10^6^ *μ*m^3^. Despite these differences, both methods provide comparable measures of linear dimensions and the identification of organisms at genus level. Furthermore, we find similar results in the range of volumes where the datasets overlap, and these distributions are also close to the distributions presented in the main text for the combined dataset.

### Sampling methods and dataset description

The measurements for the Baltic dataset were done by the HELCOM Phytoplankton Expert Group (PEG), and described in more detail by Olenina et al. (2006). The phytoplankton samples were taken in accordance with the guidelines of HELCOM (1988) as integrated samples from surface 0-10, or 0-20 m water layers, using either a rosette sampler (pooling equal water volumes from discrete depths: 1; 2,5; 5; 7,5 and 10 m) or a sampling hose. The samples were preserved with acid Lugol’s solution (Willén 1962). The inverted microscope technique (Utermöhl 1958) was used for identification of the phytoplankton species. After concentration in a sedimentation 10-, 25-, or 50-ml chamber, phytoplankton cells were measured for the further determination of species-specific shape and linear dimensions. All measurements were performed under high microscope magnification (400–945 times) using an ocular scale.

The second dataset includes the results of sampling of three to nine ecosystems per ecoregion and three locations for each system, yielding a total of 116 local sites replicated three times. Phytoplankton were collected with a 6 μm mesh plankton net equipped with a flow meter for determining filtered volume. Water samples for phytoplankton quantitative analysis were preserved with Lugol (15 mL/L of sample). Phytoplankton were examined following Utermöhl (1958). Phytoplankton were analysed by inverted microscope (Nikon T300E, Nikon Eclipse Ti) connected to a video-interactive image analysis system (L.U.C.I.A Version 4.8, Laboratory Imaging). Taxonomic identification and linear dimension measurements were performed at individual level on 400 phytoplankton cells for each sample. Overall, the data on 142 800 cells are included. The data on the dimensions of the same species were averaged for each replicate.

In both field studies, organisms’ identification was based on inverted microscopy, whenever it was not possible to reach species level, the microorganisms were identified at genus, the cells were associated with a specific geometric shape and their linear dimensions were measured. To avoid problems due to the preservation with Lugol’s solution, the samples were analysed in a short time after sampling (within few weeks).

### Cell size and shape

We classified each cell as one of 38 geometric shapes, such as spheres, cylinders, prisms, etc. (see below, and Fig. 1 for examples of phytoplankton cell shapes). We measured cell linear dimensions and calculated the surface area and volume for each cell (Hillebrand *et al*. 1999; Olenina *et al*. 2006; Vadrucci *et al*. 2007). To standardize the calculations for both databases, we have derived all formulae for surface area and volume using Maple software, yielding a list of analytic expressions for cell volume and cell surface area for each of the 38 shape types (Supplementary material for the entire list of formulae and a Maple script, which can be used as a tool for further derivations). Note that this automatic derivation allowed us to correct some of the previously published formulas.

Depending on the shape, the linear dimensions of a cell can include up to 10 measurements of different segments. To roughly characterize the cell size in 3D space, we determined 3 orthogonal dimensions of each cell, charactering the minimal, middle and maximal cell linear dimensions, which are denoted as *L*_*min*_, *L*_*mid*_ and *L*_*max*_. For simple shapes such as sphere, ellipsoid, cylinder, cone, etc., the meaning of these dimensions is clear. For some asymmetrical cells with, for instance, different horizontal dimensions at the top and bottom, we used the largest of these two dimensions, because the smallest one (or the average) does not properly describe the geometric limitations. For instance, a truncated cone is characterized by its height and two diameters: top and bottom. However, the top diameter is typically extremely small and does not reflect the cell geometry. Thus, for such shapes we used height as one dimension and the bottom diameter as the other two dimensions. For more complex shapes, consisting of few segments measured separately (e.g., half ellipsoid with a cone), we used the sum of linear dimensions of these parts as projected to each orthogonal axis.

### Measures of cell elongation

We use two characteristics of shape elongation: aspect ratio and relative surface extension. *Aspect ratio* is defined typically as the ratio between the largest and smallest orthogonal dimensions. But then we cannot distinguish between prolate (attenuated) and oblate (flattened at the poles) shapes which, according to this definition, both have aspect ratio larger than one. Therefore, it is more appropriate to define the aspect ratio *r* for prolate cells as *r = L*_*max*_/*L*_*min*_, and for oblate cells as the inverse value *r = L*_*min*_/*L*_*max*_, so that *r* < 1 for oblate shapes and *r* > 1 for prolate shapes. To distinguish between prolate and oblate cells, we use the fact that for prolate cell typically *L*_*min*_ ≈ *L*_*mid*_ < *L*_*max*_, while for oblate cells *L*_*min*_ < *L*_*mid*_ ≈ *L*_*max*_. To generalize this rule for arbitrary cell geometries, we define a cell as ‘oblate’ when its *L*_*mid*_ is closer to *L*_*min*_ than to *L*_*max*_, while for ‘prolate’ cells the opposite should be true. As cell dimensions changes for orders of magnitude, it is more convenient to use logarithmic scale for this comparison, so formally we classify a cell as *prolate*, if *L*_*mid*_ is less than the geometric mean between *L*_*max*_ and *L*_*min*_ 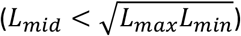 and as *oblate* in the opposite case.

In addition to ‘prolate’ and ‘oblate’, we also introduce a third, ‘compact’, shape category. We classify a cell as ‘compact’ for a certain range of *r*-values close to 1. The reason being that the aspect ratio varies over almost 4 orders of magnitude, from 0.025 to 100, and cells with a small difference in linear dimensions are closer to compact shapes than to extremely oblate or prolate forms. As the surface area increases with elongation extremely slowly, an aspect ratio of 3/2 (or 2/3) can lead to only a 2% increase in the surface area with respect to a sphere (Fig. S1). Thus, we define a cell to be ‘compact’ if 2/3 < *r* < 3/2, so that the maximal cell dimensions do not exceed the minimal dimension by more than 50%.

The second characteristic of cell elongation, the *relative surface area extension ε*, (for brevity, hereafter referred to as *surface extension*) shows the relative gain in surface area due to deviation from a spherical shape and is calculated as the ratio of the surface area *S* of a cell with a given morphology to the surface area of a sphere with the same volume *V*. Thus, 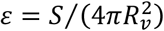, where *R*_*v*_ *=* (3*V*/(4*π*))^1/3^ is the so-called equivalent radius of a sphere with the same volume. In contrast to another measure of cell elongation, *L*_*max*_*S*/*V*, (Reynolds 1988), the surface extension directly links shape elongation at a constant volume to an increase in the surface area and therefore to potential increases in nutrient uptake or cell wall cost. Mathematically, it can also be termed the *inverse shape sphericity*, but we prefer to term it surface extension here, as it provides a more intuitive interpretation of the value. Surface extension is, in some sense, a more integrative characteristics of cell geometry than aspect ratio, as it operates with area and volume instead of linear dimensions. However, the two measures of shape elongation are related, and the logarithm of the aspect ratio changes approximately with the square root of surface extension (Fig. S1).

The minimum possible value of cell surface extension, *ε*_*min*_, is shape-specific. To find *ε*_*min*_ for a given shape (e.g. ellipses or cylinders), we need to find the combinations of shape dimensions leading to the minimal surface area for given volume. Assume that *L*_*max*_ *= αL*_*min*_ and *L*_*mid*_ *= βL*_*min*_ where *α* and *β* are some real positive numbers. For basic geometric shapes, the surface area can be expressed as 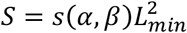 and volume as 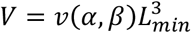, where *s*(*α, β*) and *v*(*α, β*) are shape characteristic functions that do not depend on *L*_*min*_. Then, surface extension becomes a function of only *α* and 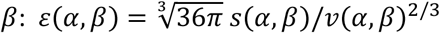. Formally, the minimal surface extension can be found as 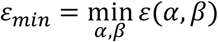 and the values 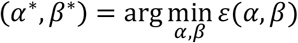 are the ratios between the linear dimensions of the specific shape with the minimal surface area. If a shape has rotational symmetry, then *α = β* and the problem becomes simpler. Solving this minimisation problem for different shape characteristic functions *s*(*α, β*) we find that for ellipses, the minimal surface extension *ε*_*min*_ *=* 1 is achieved when all semi-axes are equal, that is, if the ellipse is a sphere. For a cylinder *ε*_*min*_ *=* (3/2)^1/3^ *=* 1.14, when its height equals the diameter; for a parallelogram or prism on a rectangular base *ε*_*min*_ *=* (6/*π*)^1/3^ ≈ 1.24 (when it is a cube). In all these cases *α*^∗^ *= β*^∗^ *=* 1, which means that the minimal surface area for these shapes is achieved when all linear dimensions are equal.

### Statistical analysis

For the present analysis, to reduce variability in cell sizes, the data was averaged for each genus and local site. We chose averaging at the genus rather than species level, because not all cells were identified at the species level.

For making histograms, we binned the data using a linear scale for binning surface extension and logarithmic scale for binning volume. For distribution of genus richness over cell volume, we used 30 bins in the range [0.1, 10^9^], for distribution of richness over surface extension we used 40 bins in the range [1, 5], and for bivariate distributions of richness, the data was binned into 10 volume classes and 15 surface extension classes using uniform ranges. To fit nonlinear distributions, we used MATAB function fitnlm, which uses the Levenberg-Marquardt nonlinear least squares algorithm.

### Modelling effects of geometric constraints

To make a theoretical prediction of a potential variation in cell elongation for cells of different volume we calculate the surface area and volume of an ensemble of 50,000 ellipsoidal cells whose dimensions are constrained according to two scenarios. In the first scenario we randomly draw linear dimensions of the ellipses from a log-uniform distribution in the range 1 ≤ *L*_*min*_, *L*_*mid*_, *L*_*max*_ ≤ 1000 *μm*. In the second scenarios we additionally assume that the aspect ratio *r* is constrained by a sigmoidal function of cell volume to not exceed the maximal observed values of *r* in data, see Results and Table 1.

**Table 1.**
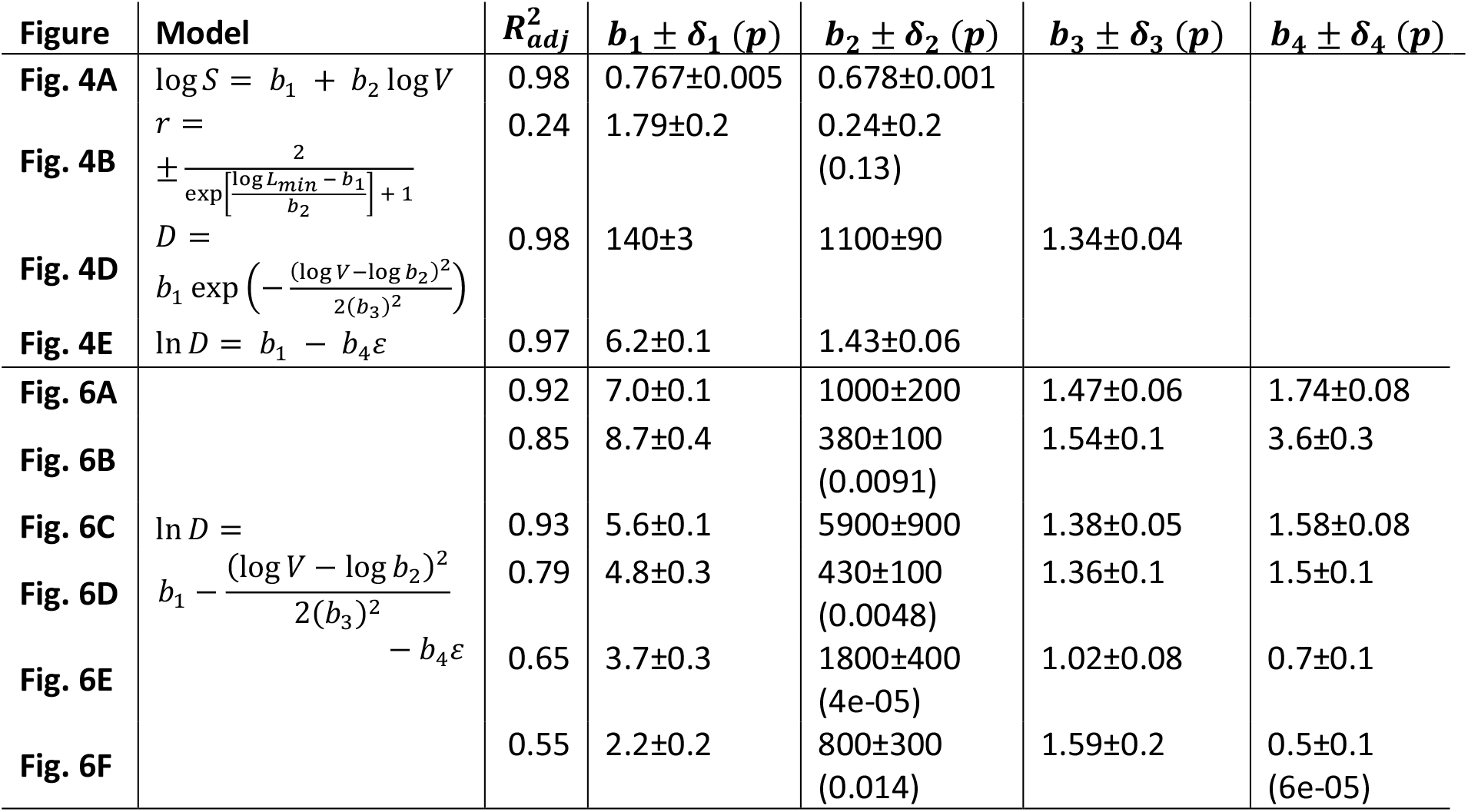
Fitting parameters for Fig.4 and Fig.5. Parameter values *b*_*i*_ are specified with standard error *δ*_*i*_ and p-value in brackets (only when *p* > 10^−5^). Fitting in Fig. 4B is done to the outer hull of the data points.

### Intracellular diffusion constraints

Linear dimensions of phytoplankton cells in our database are less than 1000 *μ*m. One possible explanation is that the maximal cell size can be constrained by the distance of intracellular diffusion during one cell life cycle (Gallet *et al*. 2017). The mean diffusive displacement of particles in 3D space equals 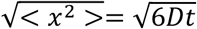, where D is the diffusion coefficient and *t* time interval. The diffusion coefficient of proteins in cytoplasm of bacteria, *Escherichia coli*, ranges from 0.4 to 7 *μm*^2^/*s* (Kumar *et al*. 2010). Diffusion rates in cytoplasm measured by Milo & Phillips (2020) lay also in this range. According with this data, the mean diffusive displacement in the cell cytoplasm during one day (a typical reproduction time scale for phytoplankton) should range from 455 to 1900 μm.

## Results

### Cell shapes with most taxonomic diversity across phyla

Our data contain 402 phytoplankton genera of various shape and size (Fig. 1). The taxonomic diversity of genera changes across phyla and cell shape type, with some shapes being much more common and occurring among many genera. The shapes exhibiting the highest taxonomic diversity depended on the phylum (Fig. 2A). There is a clear difference in which shapes are prevalent among Bacillariophyta (diatoms) vs. other phyla. While most diatom genera are cylindrical, prismatic and rhomboid, among other phyla, the highest taxonomic diversity is observed for ellipsoidal cells, with conical or more complex shapes occurring among only a few species or genera. In our database, 46% of the genera are prolate, 38% compact and only 16% oblate. However, this ratio depends on the cell geometric shape (Fig. 2B). While more than half of the genera with elliptic cells have a compact shape, other shapes have more than half of genera with prolate cells.

**Fig. 2.**
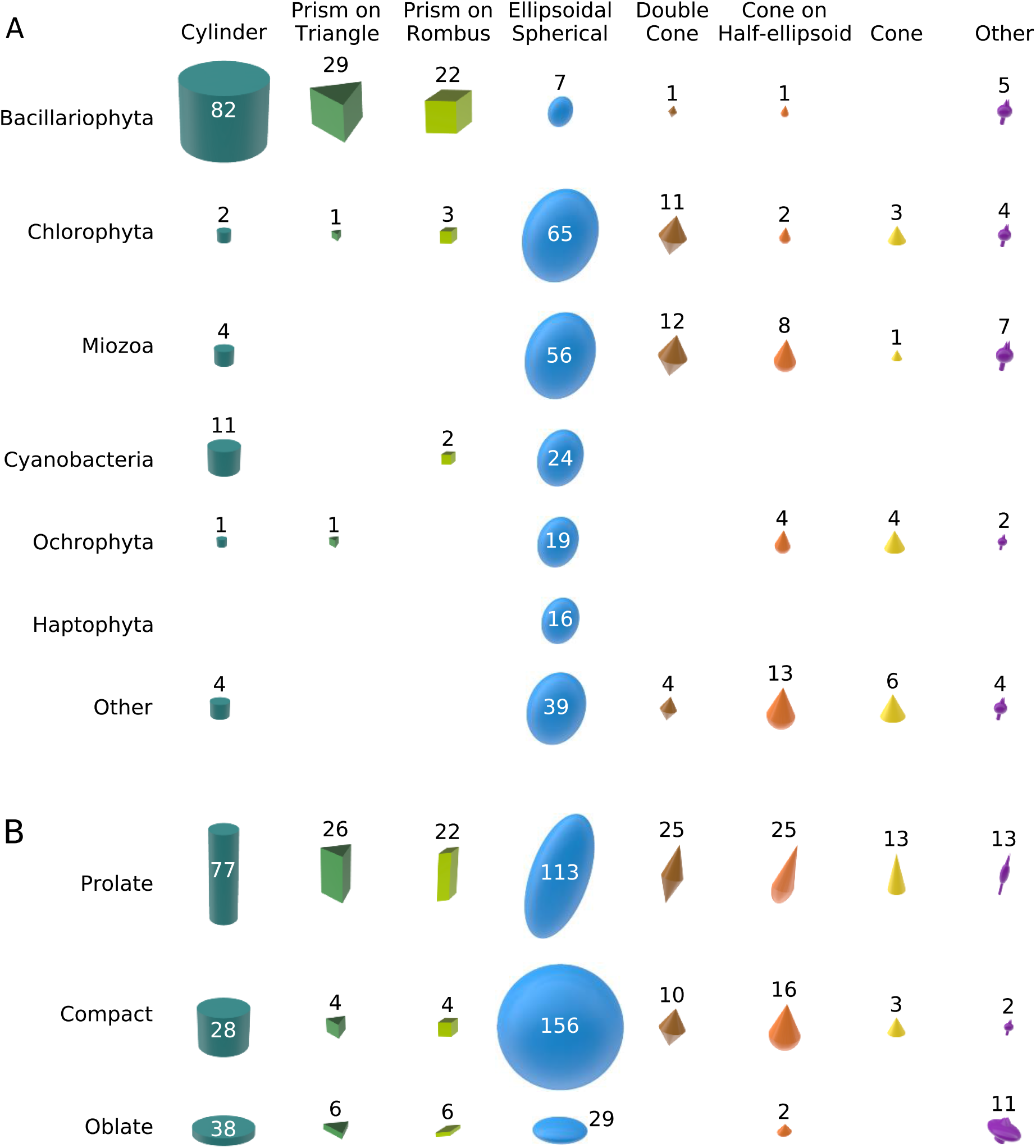
Diversity distribution of various shape types (columns) across phyla (A, rows) and across cell shape elongation (B, rows). The area of each figure is proportional to the number of genera (shown next to or within it). See Fig. 3 for detailed analysis. Note that if a genus includes cells with different shapes, this genus is included in several cells in panel **(A)**. Thus, the total number of genera in panel **(A)** exceeds the total number of genera in our database. The same applies to panel **(B)**, as cells of the same genus can be assigned to more than one elongation and shape type.

Shape, elongation and phyla are interrelated (Fig. 3). For most phyla (except of Bacillariophyta, Miozoa, Haptophyta, Charophyta, and Euglenozoa) the largest taxonomic diversity is observed in the classes of prolate or compact cells (around 40-50% of genera in each class) with oblate cells displaying relatively low diversity (< 10% of genera). By contrast, most diatom genera have either prolate (60% of genera) or oblate shape (25%) with only 15% of genera being compact. For Haptophyta, Charophyta, and Euglenozoa we find a similar distribution with a small fraction of compact cell genera and a relatively large faction of oblate and prolate cell genera. Dinoflagellates (Miozoa) have also a relatively large fraction of genera with oblate cells (18%) and almost equal factions of compact and prolate cells (around 40% of genera in each group).

**Fig. 3.**
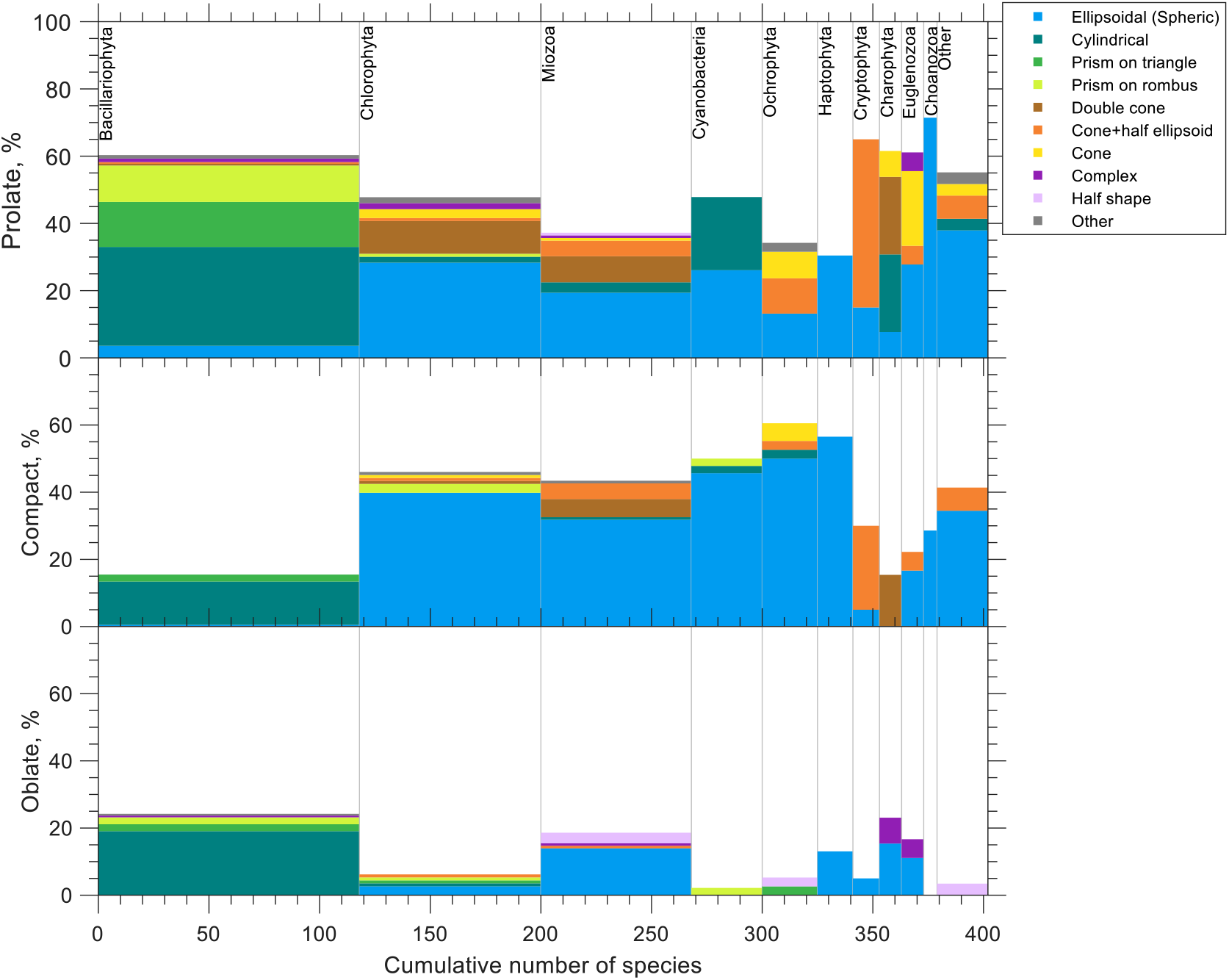
Diversity of phytoplankton genera across cell shapes (colour coded) and shape elongation (top, middle and bottom panel) for different phyla (columns). Most of compact and prolate cells have cylindrical or prismatic shape in Bacillariophyta, conic shapes in Cryptophyta and Charophyta, and ellipsoidal shapes in the other phyla. Oblate cells are present in Bacillariophyta, Miozoa and Haptophyta, while for the other phyla their frequency is less than 10%, in particular oblate cells absent in cyanobacteria, Ochrophyta and Cryptophyta. Most of cylindrical and prismatic species belong to Bacillariophyta. Bacillariophyta almost do not contain ellipsoids which have a large fraction in the other phyla. See main text for further detail.

Prismatic and cylindrical shapes are common in diatoms and cylindrical prolate shapes in Cyanobacteria and Chlorophyta. In other phyla, more than 60% of genera are elliptic with a significant fraction (∼20%) of conic shapes. Half shapes, such as half-spheres or half-cones are relatively rare and typically are found in oblate forms of Miozoa, Ochrophyta and some other phyla. Complex shapes (such as an ellipse with cones or cylinders) comprise 10-20% of genera in Charophyta and Euglenozoa but are extremely rare in other phyla.

### Cell elongation and volume

Cell volumes in our database span almost 10 orders of magnitude, from 0.065 *μ*m^3^ for the cyanobacterium *Merismopedia* to 5 · 10^8^ *μ*m^3^ for Dinophyceae*’*s *Noctiluca*. In contrast to previous studies (Lewis 1976; Niklas 2000), our analysis shows that cell surface area increases with volume approximately to the power of 2/3 (Fig. 4A), indicating that cell dimensions scale on average isometrically with volume, and there is no evidence for more shape elongation with increasing volume.

**Fig. 4.**
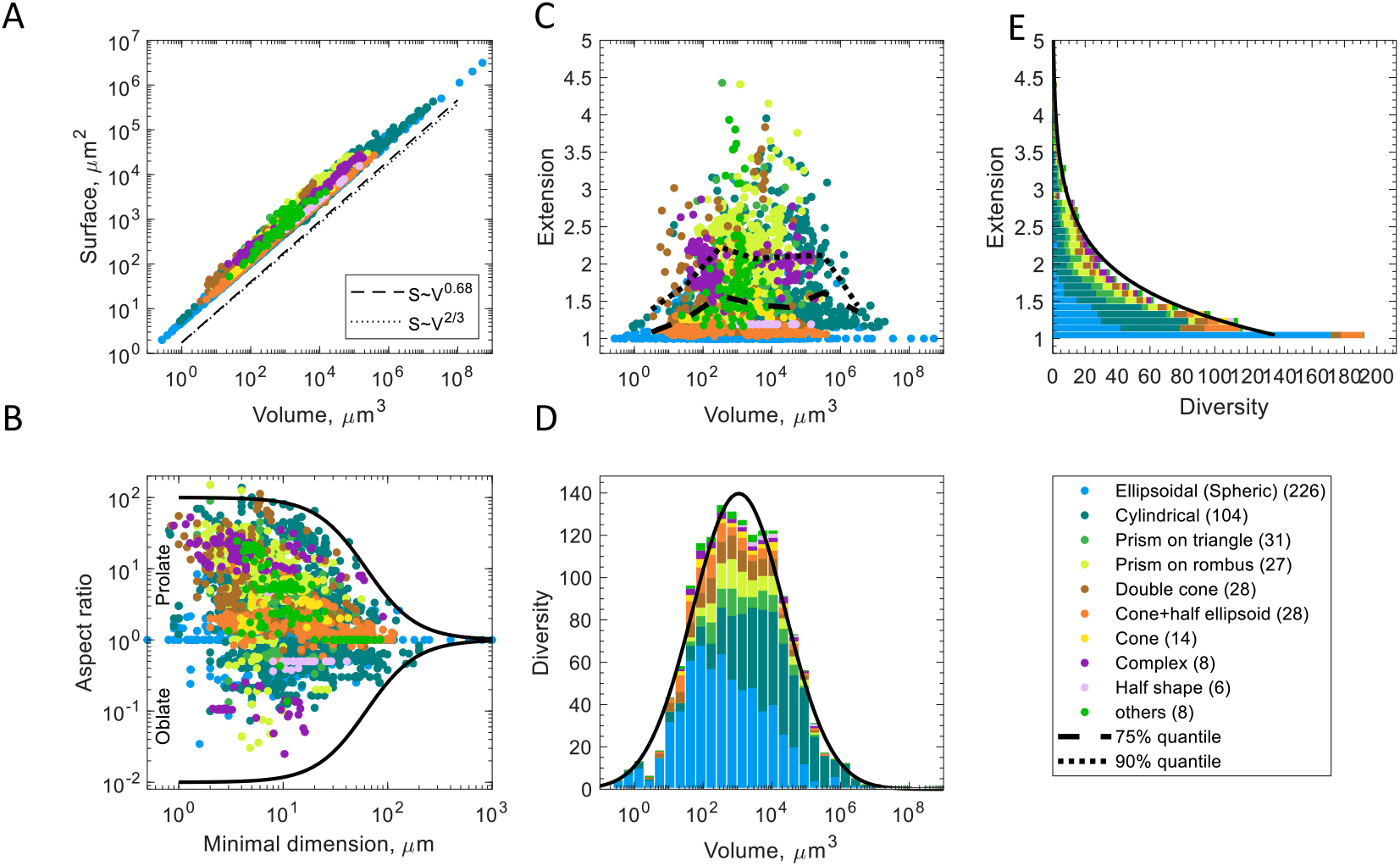
Geometry of unicellular phytoplankton for various cell shape types. (A) Surface area as a function of cell volume. The dashed, and dotted, lines show the slope of a power law fit, and a scaling with the power of 2/3, respectively. (B) Aspect ratio, *r*, as a function of minimal cell dimension. The solid line shows a fitted sigmoidal function to the upper boundary of | log *r* | (black solid line). (C) Surface relative extension as a function of cell volume. The dotted and dashed black lines show 75% and 90% quantiles. (D) Distribution of taxonomic diversity as a function of cell volume. The black line shows a fitted Gaussian function. (E) Distribution of taxonomic diversity over cell surface extension (note the interchanged axes). The black line shows a fitted exponential function. The legend depicts the colour coding for different shape types, with the number of genera for each shape type given in parenthesis. See Table 1 for fitting parameters.

The extent of cell elongation strongly varies with cell volume, following a hump-shaped distribution (Fig. 4C). Cell surface extension exhibits the largest variation at intermediate cell volumes (between 10^3^ − 10^4^ *μm*^3^), in this range covering an enormous variety of shapes including compact, oblate, and elongated forms, with maximal surface areas exceeding that of a sphere by up to 5-fold (Fig. 4C). By contrast, for cells of very small or large volume, surface extension approaches its minimum values, implying that these cells have a compact shape minimizing their surface area. The hump-shaped pattern is also seen in the 75% and 90% quantiles (Fig. 4C, black dashed and dotted lines), confirming that this is not a sampling artifact related to a smaller number of the large and small volume cells compared to the number of intermediate volume cells. Also, the aspect ratio varies the most for intermediate cell volumes, spanning from 1/40 for oblate cells to 100/1 for prolate cells, confirming that our dataset includes the entire spectrum from oblate to extremely elongated shapes (Fig. 5A). This pattern holds across different trophic guilds (autotrophic, mixotrophic or heterotrophic); however, the maximum cell elongation is reached only by autotrophs, while in heterotrophs and mixotrophs the maximum aspect ratio is 10 and the maximum surface extension is 2 (Fig. S2), likely because these two groups need to swim actively and have a more complex internal organization (Kiørboe 2008).

**Fig. 5.**
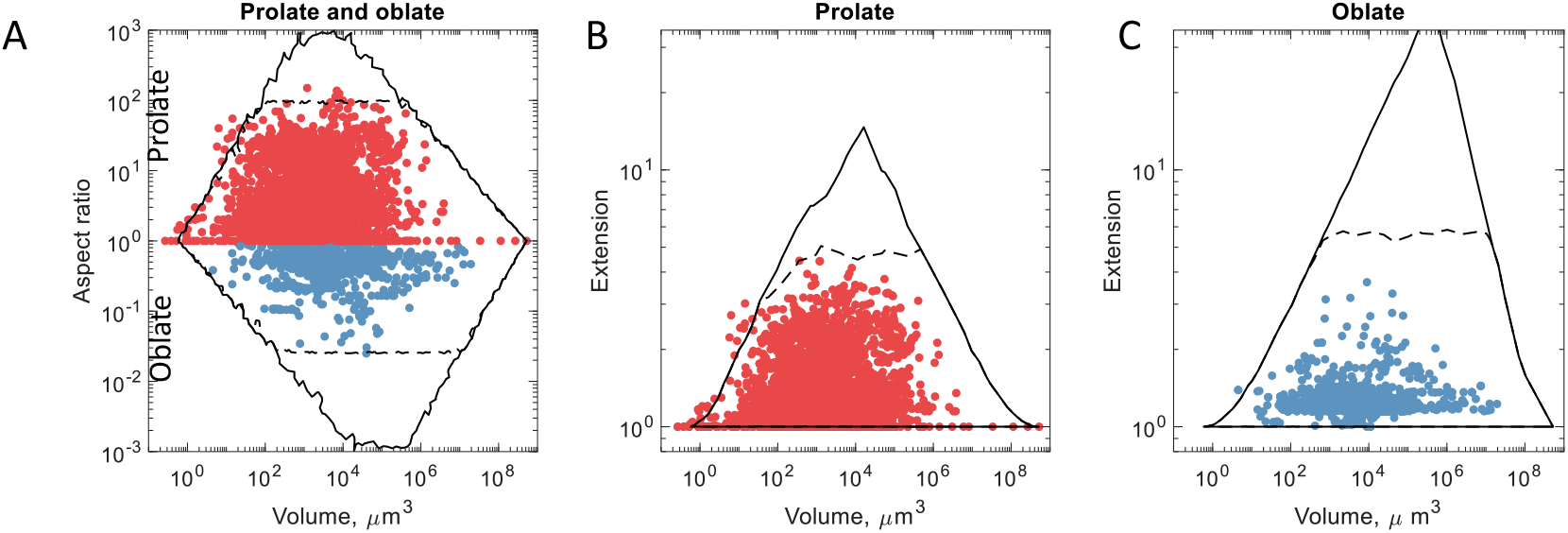
Aspect ratio and surface extension of oblate and prolate cells compared to model predictions. (A) Comparison of the aspect ratio of prolate (red circulars) and oblate (blue circulars) cells with outer hulls for volume and aspect ratio of 50,000 ellipsoids with dimensions randomly chosen according to the first scenario (black line) and second scenario (black dashed line). See Methods for the description of scenarios. (B, C) the same for combinations of volume and surface extension for prolate (B) and oblate (C) cells.

### Phytoplankton diversity distributions

Taxonomic diversity, *D*, measured here as richness of genera, depends both on cell volume and surface extension. The distribution of diversity across cell volume follows a lognormal function of cell volume with a peak of diversity at *V*_0_ *=* 1100 ± 90 *μm*^3^ (Fig. 4D, 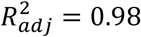). When data are binned over surface extension, the distribution decreases exponentially with shape surface extension as *D*∼*e*^−1.43*ε*^ (Fig. 4E, 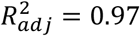). Both relationships depend on cell shape (Fig. S3, S4). The ellipsoidal cells have the highest genus diversity at the smallest volume compared to other shapes (*V*_0_ *=* 330 ± 40 *μm*^3^, 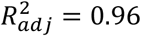) and the fastest rate of diversity decrease with surface extension (*D*∼*e*^−2.4*ε*^, 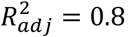), with 54% of the genera exceeding the surface area of a sphere by less than 10%. In contrast, for cylindrical cells (mainly diatoms), diversity peaks at the largest volume compared to other shapes (*V*_0_ *=* 8,700 ± 800 *μm*^3^, 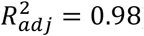) and declines more slowly with surface extension (*D*∼*e*^−1.4*ε*^, 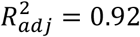). There is a comparable effect of surface extension on diversity for conic shapes (*D*∼*e*^−1.2*ε*^, 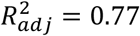). The effect is weaker for prismatic (*D*∼*e*^−0.95*ε*^, 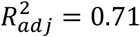) and complex shapes (*D*∼*e*^−0.75*ε*^, 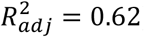), noting that both prismatic and complex shapes occur mainly in diatoms. The secondary peaks of diversity occur at *ε* between 1.5 and 3 for prismatic and complex shapes. The weaker correlation of diversity with cell elongation for complex shapes could also be caused by the fact that representing complex shapes requires more parameters than just simple composites, such as aspect ratio or surface extension.

Thus, both cell volume and surface extension correlate with taxonomic diversity. Assuming that volume and surface extension affect species fitness independently of each other, we can approximate the diversity distribution as a product of a lognormal function of volume and a decreasing exponential function of surface extension

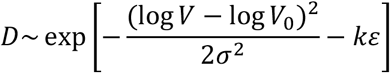

As shown in Fig. 6, this function describes the dependence of diversity on both cell volume and surface extension remarkably well, with *V*_0_ *=* 1,000 ± 200 *μm*^3^ (mean volume), σ *=* 1.74 ± 0.08 (variance of the logarithm of volume) and *k =* 1.47 ± 0.06 (the rate of exponential decrease of diversity with surface extension), explaining 92% of the variation of phytoplankton diversity for the entire dataset. We obtain nearly identical distributions both for the combined data set and for each of the two data sources separately (Fig. S5).

**Fig. 6.**
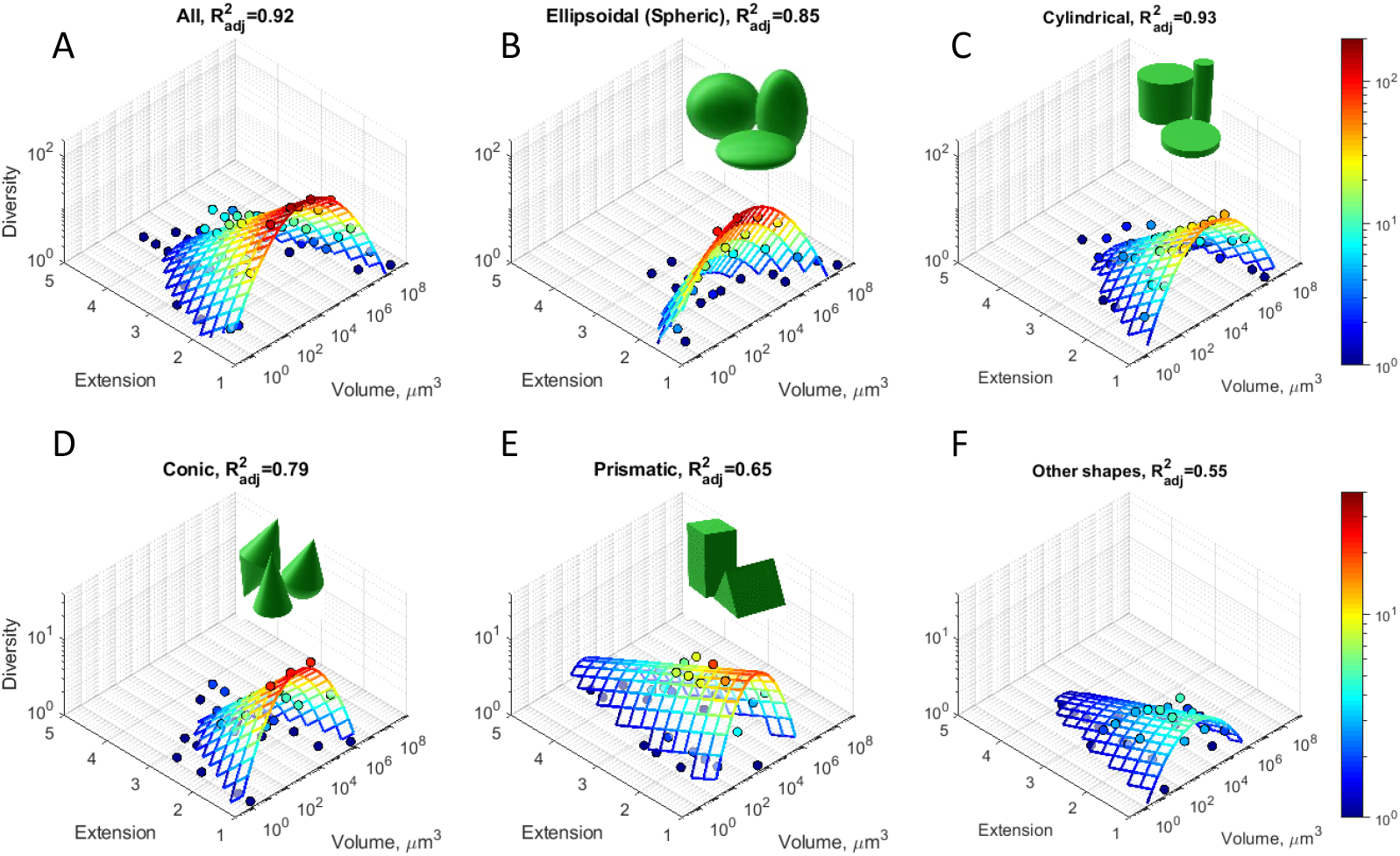
Diversity distribution of unicellular phytoplankton. (A-F) Bivariate histograms of taxonomic diversity, *D*, as a function of surface extension, *ε*, and logarithm of cell volume, *V*, (dots), aggregated over all shape types (A) and for different shape types (B-F). Note that due to intraspecific and intragenus variability cells of the same genera can contribute to diversity in different bins. The mesh (solid lines) shows a fit by the function In *D = a* −(Iog *V* − Iog *V*_0_)^2^/(2 σ^2^) − *kε*, weighted with diversity. The colours indicate taxonomic diversity from *D =* 1 (blue) to *D =* 200 (red) in A-C and to *D =* 40 in D-F. See Table 1 for regression results, and Fig. S6 for comparison between predicted and observed diversity.

Across different shapes, the fitted parameters have the same trends as above: the best match is obtained for ellipsoidal, cylindrical and conic shapes (Fig. 6B-D), with a poorer fit for prismatic and other shape types (Fig. 6E-F). A comparison of the predicted and the observed diversity shows that there is an unbiased fit for all shapes combined, and also in the group of ellipsoidal, cylindrical and conic shapes (Fig. S6A-D). However, the fit for prismatic and other shapes overestimates taxonomic diversity for the volumes and surface extensions where the observed diversity is low (Fig. S6E-F). Note that the correlations in Fig. 6, derived for all shapes combined (*R*^2^ *=* 0.92), are higher than those obtained for some specific shapes. This is related to the fact that different shape classes exhibit diversity peaks at slightly different values of surface extension (e.g., ellipsoidal cells exhibit maximal diversity at *ε =* 1, while that of prismatic cells peaks at *ε* ≈ 1,5). This separation reduces the quality of fit for specific shape types but does not play a role when we consider all shapes together (compare Fig. S4A with Fig. S4B-F).

The predictions for taxonomic diversity based on aspect ratio are, on average, less strong than those based on surface extension. Regression analysis of the diversity distribution across volume and aspect ratio gives 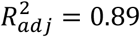 for all data and 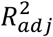 ranging from 0.23 to 0.86 for specific shapes (Fig. 7). The reduced 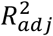 values compared to the fitting based on surface extension probably occur because of a more complicated functional dependence of diversity on aspect ratio (Fig. S7). For instance, for ellipsoidal prolate shapes diversity monotonically decreases with aspect ratio but shows a peak for oblate shapes at *r* ≈ 1/2. For cylinders, the picture is even more complicated with two peaks of diversity at *r* ≈ 3 and 1/3.

**Fig. 7.**
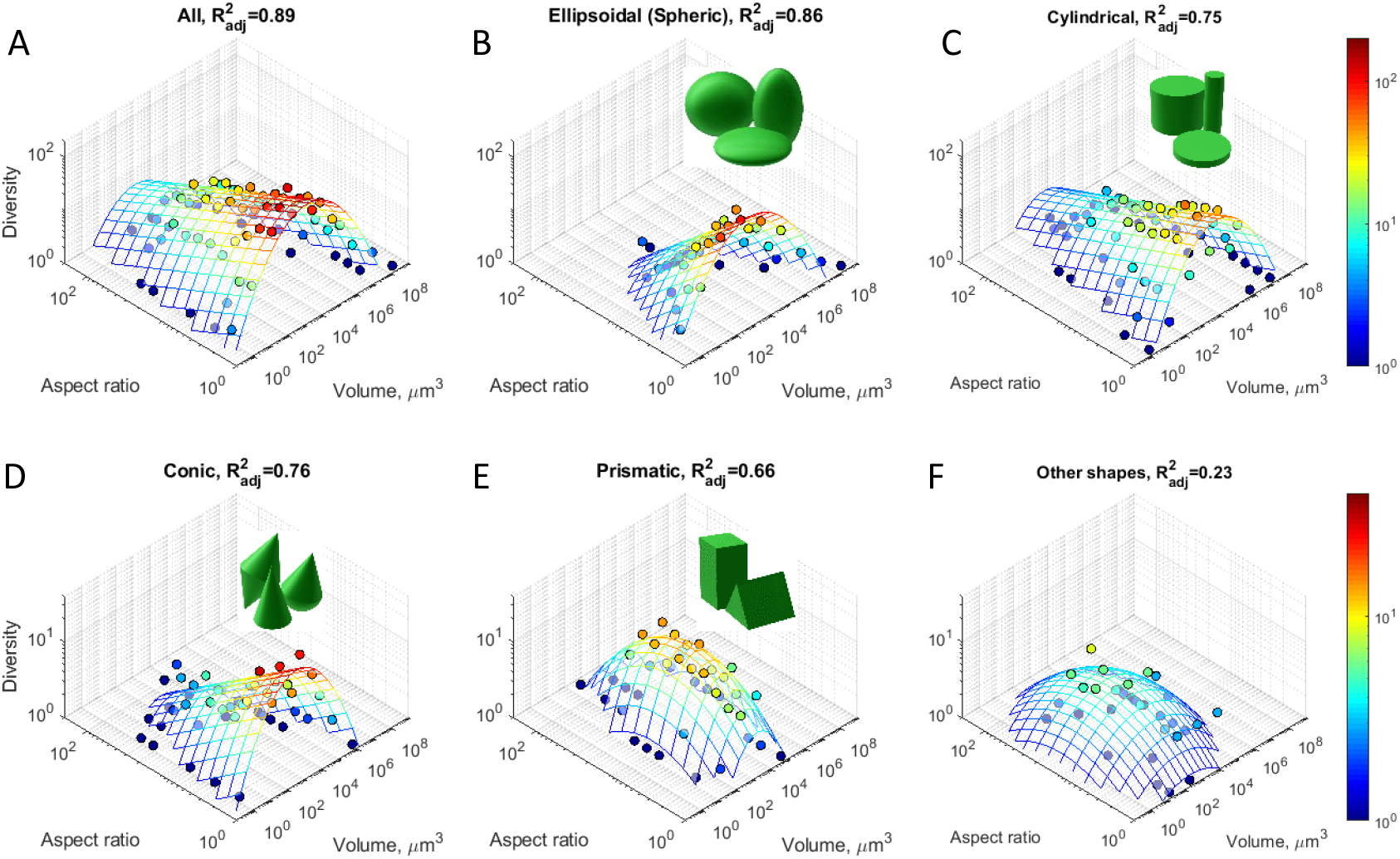
Diversity distribution of unicellular phytoplankton. Bivariate histogram of taxonomic diversity as a function of aspect ratio and volume. To reduce the number of fitting parameters the aspect ratio here is measured as *L*_*max*_/*L*_*min*_, so that no distinction between prolate and oblate cells has been made. Note that due to intraspecific and intragenus variability cells of the same genera can contribute to diversity in different bins. To provide a better fit for prismatic and other shapes (**E, F**), where diversity peaks at intermediate values of the aspect ratio, we also assumed a log-normal dependence on the aspect ratio. See Table S1 for the results of regression analysis.

The difference between how diversity depends on surface extension vs. aspect ratio likely stems from the nonlinear relationships between these parameters (Fig. S1). The logarithm of aspect ratio changes approximately as 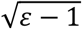, implying an extremely high rate of change of the aspect ratio with *ε* for compact shapes, and a much smaller rate for elongated shapes. Consequently, projecting diversity onto the surface extension axis results in an exponential decrease, while projecting it on the aspect ratio axis results in a bimodal distribution with a local minimum of shape diversity for *r =* 1 (Fig. S1B,C). However, these projections show only a part of the entire picture. As shown in the bivariate plot (Fig. S1A), diversity peaks for spherical cells (both surface extension and aspect ratio of around 1) and then decreases with further deviation from this shape towards prolate or oblate forms. This decrease is asymmetric and occurs faster for oblate shapes.

## Discussion

Our analysis of phytoplankton cell sizes and shapes within the extensive dataset we assembled reveals several novel patterns, shedding light on the morphological and taxonomic diversity in this globally important group of marine microbes. We show that there is an interplay between different cell sizes and shapes, where the cells of intermediate volumes can have very diverse shapes and range from oblate and to extremely prolate forms, while cells of both large and small volumes are compact (mostly spherical). At the same time, spherical shapes exhibit the largest variation of cell volumes. Finally, taxonomic diversity has a peak for compact cells of intermediate volume and decreases exponentially with cell surface extension for attenuated and flattened cells.

### Diversity changes with cell volume and surface extension

Our study shows that cell surface extension, in addition to cell size, correlates with diversity, with the two traits together explaining up to 92% of its variance. The diversity distribution follows a lognormal function of volume, and decreases exponentially with cell surface extension. This pattern is likely to be universal, as we have obtained similar biodiversity distributions both for the combined dataset and separately for each of the two datasets (Baltic Sea and 6 ecoregions of the world’s oceans) (Fig. S5). This suggests that these traits may be important drivers of diversity. As diversity typically increases with abundance (Siemann *et al*. 1996), we hypothesize that species with compact cells of intermediate volume are the most adapted among unicellular plankton species for survival in permanently changing water conditions.

Thus, for all phyla, except for prismatic and complex shapes (mainly diatoms), a reduction of cell surface area is likely an advantageous strategy, which leads to greater diversification rates and higher diversity of compact cells compared to elongated cells in each cell volume class. Reducing cell surface area likely reduces the cost of cell walls and makes a cell less vulnerable to predators. However, non-spherical elongated shapes can be cheaper and advantageous for species with rigid cell walls, such as diatoms (Martin-Jézéquel *et al*. 2000; Monteiro *et al*. 2016). This can explain why, for prismatic and complex shapes (mainly diatoms), we observe secondary peaks in richness for elongated shapes, resulting in significant diversity of diatom shapes and taxa across a wide range of cell elongation. In these taxa, cell elongation can have a nonmonotonic effect on cell fitness, such that both compact and elongated cells can have high diversity (Grover 1989). This suggests that the appearance of silica cell walls in diatoms is a major evolutionary innovation that allows diatoms to achieve an unusually large shape diversity, which may have contributed to the ecological success of this group (Nelson *et al*. 1995; Malviya *et al*. 2016).

### Elongation and linear dimensions

To what extent can these patterns in biodiversity be explained by constraints on cell dimensions? Linear cell dimensions in our data range from 0.5 *μ*m to 1,000 *μ*m (Fig. 4B). The minimum cell size is likely constrained by the size of organelles; for instance, for autotrophs the minimum chloroplast size equals 1 *μm*(Raven 1998; Li *et al*. 2013). The maximum cell size of unicellular organism can be constrained by diffusive scale, mechanical stability, or metabolic optimality. Firstly, the maximal cell size can be constrained by the intracellular diffusion rate. For instance, to homogenously distribute molecules within a cell, its size should not exceed the mean diffusive displacement of molecules in cell cytoplasm during one life cycle. A simple calculation gives a range from 455 to 1900 μm (Methods). More detailed calculations show that cell size can effect intracellular diffusion and therefore metabolic rates; thus, larger bacterial cell volumes become possible likely due to a reduction of molecular transport time inside the cell (Gallet *et al*. 2017). Second, to avoid mechanical damage in moving water, a cell should be smaller than the smallest eddies which have a size of around 200 *μm*(Reynolds 1988). Last but not least, differences in scaling of metabolic rates for different resource strategies might make large unicellular organisms suboptimal compared to multicellular organisms (Andersen *et al*. 2016). Metabolic rates can become limited by the nutrient uptake rate as the surface to volume ratio decreases with increasing volume (Reynolds 2006).

Given that the cell dimensions are constrained within a fixed range, minimal (or maximal) cell volume can only be realized in a compact geometry when all three linear dimensions are equal to the minimal (or maximal) possible value within this range, while the largest variation of cell shape will be possible for intermediate volumes. To check if this geometric constraint can explain the patterns shown in Fig. 4C and 5, we calculated surface area and volume for an ensemble of elliptical cells with dimensions randomly drawn from the range of 1 to 1000 *μm*(Methods). Similar to the empirical data, the smallest and largest cells in this ensemble are compact, while cells of intermediate volumes have a diverse geometry (Fig. 5, solid line). However, this approach overestimates the maximal possible aspect ratio (ranges from 10^−3^ to 10^3^) and surface extension (yielding values of *ε* > 10 for prolate cells and *ε* > 30 for oblate cells). In a second scenario, we assumed additionally that cell aspect ratios do not exceed the maximal observed values of *r* (the envelope shown as black line in Fig. 4B). As the longest linear cell dimension *L*_*max*_ < 1000 *μm*, the allowed range of *r* reduces with increasing the shortest cell dimension *L*_*min*_, so that *r* approaches 1 when *L*_*min*_ approaches 1,000 *μ*m. This constraint may reflect an additional limitation due to mechanical instability (Reynolds 1988), material transport requirements within a cell, or reduced predator defence experienced by extremely prolate or oblate cells. After imposing this constraint on the range of aspect ratios, the model and the data agree well for prolate cells, but the theoretical model still overestimates the potential surface extension for oblate cells of large volumes (Fig. 5, dashed line).

This suggests that there may be some additional constraints that prevent the evolution of extremely wide oblate cells with a large volume. In particular, even having the same aspect ratio, oblate and prolate cells might differ in their sinking velocity (Padisák *et al*. 2003) and nutrient uptake rate (Karp-Boss & Boss 2016). A flat thin disc has a larger surface area than an elongated cylinder with the same volume and aspect ratio. However, an increased *S*/*V* ratio will not necessarily increase the flow of nutrients, because it also depends on the cell shape. Nutrient concentrations are depleted in the extracellular environment around the cell boundary and the diffusion rate of new nutrients towards the cell wall declines as the cell curvature decreases, e.g., when cell radius increases (Kiørboe 2008). As a result, elongated cells can provide a larger nutrient flow compared to oblate cells with the same surface area and volume (Karp-Boss & Boss 2016). This mechanism might explain the greater richness of elongated shapes compared to oblate ones (Fig. 3), as well as the fact that the maximum observed surface extension for oblate shapes is even smaller than that for elongated shapes (Fig.5 B, C).

We find that, on average, surface area increases approximately isometrically with cell volume (Fig. 4A). Thus, the surface to volume ratio decreases approximately as *V*^−1/3^. In particular, when cell volume increases from 1 *μ*m^3^ to 10^7^ *μ*m^3^, *S*/*V* decreases for almost 200-fold (from 5.8 *μ*m^−1^ to 0.03 *μ*m^−1^). Cell attenuation or “flattening” can offset only a small part of this decrease, because the most attenuated shapes in our database have surface extension less than 5, and even for a hypothetical extremely elongated cell with aspect ratio of 10,000 this factor is less than 10 (Berg 1993). Furthermore, only cells of intermediate volume can take extremely elongated or oblate forms, while the elongation of large cells is constrained by the maximal values of *L*_*max*_. Thus, large cells should have additional mechanisms compensating their low *S*/*V* ratio such as increasing the density of nutrient accommodating sites (Aksnes & Cao 2011), the presence of vacuoles (Kiørboe 2008; Litchman *et al*. 2009; Tambi *et al*. 2009; Kerimoglu *et al*. 2012) and buoyancy regulation (Reynolds 2006).

### Cell elongation and environment

The surprisingly good prediction of global taxonomic richness of marine plankton by cell volume and surface relative extension implies either a fundamental metabolic relationship between these parameters and speciation rates or a specific global distribution of niches favouring oblate (and prolate) shapes in competition with compact shapes, as the environment can select certain cell morphologies (Kruk & Segura 2012; Charalampous *et al*. 2018). In particular, very elongated shapes occur mostly in deep waters (Reynolds 1988). This can be explained by the fact that elongated shape optimizes packing of chloroplasts along the cell surface and increases light harvesting (O’Farrell *et al*. 2007). Our research provides an additional argument for the hypothesis that the dominance of elongated cells in deep water is due to the fact that these waters are also rich in the nutrients needed for walls of such cells (Reynolds 2006). Light and nutrients typically have opposing gradients (Klausmeier and Litchman, 2001, Ryabov and Blasius, 2011), so that low-light and high-nutrient conditions often co-occur in deep waters. Therefore, one can expect an increase of cell elongation with depth in poorly mixed waters. Even in well-mixed waters, Naselli-Flores and Barone (2007, 2011) found that cell elongation increases with decreasing light intensity only when species associated with high nutrient concentrations dominate.

A link between phytoplankton diversity and morphology has not been explored in detail before; and previous studies on the topic did not find a consistent pattern. In particular, local species richness showed either a hump-shaped function, was independent of cell volume (Cermeño & Figueiras 2008), or decreased as a power function of volume (Ignatiades 2017). There may be several explanations for the discrepancy between our and these previous results. Firstly, unlike previous studies, our focus on cell surface extension as an important parameter allows separation of its effects from those of cell volume. Second, our study includes a much wider range of cell volumes and more species. Lastly, it includes samples from world’s ocean ecosystems of various typology and in different times of the year, so this global pattern may be different from the local patterns influenced by specific environmental conditions, such as nutrient or light levels, grazing, species sorting or mass effects.

### Open questions

Our findings show that taxonomic richness correlates not only with cell size but also with cell shape, opening new avenues of biodiversity research. For example, environmental factors could, through affecting cell shape and size differently, drive shape-size distributions of phytoplankton assemblages, and thereby biodiversity. In particular, temperature, salinity and nutrients often change cell volume and species composition and may, thus, alter diversity (Agawin *et al*. 2000; Acevedo-Trejos *et al*. 2013). These effects would be important to investigate in the context of both periodic seasonal changes and rapid ongoing environmental change. Indirect changes in diversity and community composition can be caused by grazing, through its differential effect on cells of various shapes and sizes or by environmental factors through a potential link between cell elongation and a trade-off between generalist and specialist strategies or other ecological trade-offs. Finally, since many phytoplankton genera are present in the natural environment as colonies or chains, colony size, shape and the geometry of chains formation might also become important evolutionary factors leading to species dominance or high speciation rates. Answering these questions would help us further understand the ecological and evolutionary constraints on phytoplankton diversity in the ocean.

Our study focuses on unicellular organisms, leaving the question open as to how body shape affects evolutionary success in higher life forms. For instance, how do leaf and petal shape affect plant diversity, and which shapes result in the highest taxonomic richness? Another important question is regarding mechanistic drivers of shape optimality: is shape mainly influenced by light or nutrient harvesting, mechanical instability, maximization or minimization of surface area? Finally, questions remain regarding the distribution of species with suboptimal shapes. Our study indicates that diversity decreases with cell elongation, but this decrease is not abrupt, and elongated species constitute an essential fraction in any shape type (Fig. 2B). Thus, the rate of this decrease and its dependence on the environmental conditions may provide vital information about the distribution of ecological niches suitable for species of various morphology.

## Supporting information

38 geometric shapes

## Acknowledgements

We thank H. Hillebrand for useful comments on the manuscript and Michael Guiry for providing data on phylogenetic classification. A.R. acknowledges funding by the Ministry of Science and Culture, State of Lower Saxony project POSER, funding by HIFMB a collaboration between the Alfred-Wegener-Institute, Helmholtz-Center for Polar and Marine Research, and the Carl-von-Ossietzky University Oldenburg, initially funded by the Ministry for Science and Culture of Lower Saxony (MWK) and the Volkswagen Foundation through the “Niedersächsisches Vorab” grant program (grant number ZN3285). E.L. was in part supported by the sabbatical fellowship from the German Centre of Integrative Biodiversity (iDiv). A.B. acknowledges funding by the Puglia Region, through the strategic project PS126 ‘PhytoBioImaging’. O.K and. B.B. acknowledge funding by the German Research Foundation, DFG through the projects KE 1970/1-1 and BL 772/6-1 respectively, within the Priority Programme 1704 DynaTrait. OK was additionally supported by DFG through the project KE 1970/2-1.

## Author contributions

A.R. designed the research and performed the analysis with contributions by O.K; A.R. and O.K. calculated cell surface and volume; A.R. wrote the manuscript with contribution from O.K., B.B., E.L., I.O., L.R.; I.O. and L.R. described methods. I.O., L.R. A.B., E.S. provided data.

Correspondence and requests for materials should be addressed to A.R.

### Data availability

Data on Baltic sea are publicly available under http://ices.dk/data/Documents/ENV/, (ICES CEIM), Data on the global ecosystems are available under https://dataportal.lifewatchitaly.eu/data, (LifeWatch ERIC). The original and compiled datasets are also available on DataDryad.org under https://doi.org/10.5061/dryad.r7sqv9sb6.

### Code availability

Data analysis was implemented in MATLAB R2019. The source code for calculating volume and surface area of shapes is publicly available on GitHub (https://github.com/AlexRyabov/Cell-shape).

## References

Acevedo-Trejos, E., Brandt, G., Merico, A. & Smith, S.L. (2013). Biogeographical patterns of phytoplankton community size structure in the oceans. Glob. Ecol. Biogeogr., 22, 1060– 1070.

Agawin, N.S.R., Duarte, C.M. & Agustí, S. (2000). Nutrient and temperature control of the contribution of picoplankton to phytoplankton biomass and production. Limnol. Oceanogr., 45, 591–600.

Aksnes, D.L. & Cao, F.J. (2011). Inherent and apparent traits in microbial nutrient uptake. Mar. Ecol. Prog. Ser., 440, 41–51.

Albert, J.S. & Johnson, D.M. (2012). Diversity and evolution of body size in fishes. Evol. Biol., 39, 324– 340.

Alós, J., Palmer, M., Linde-Medina, M. & Arlinghaus, R. (2014). Consistent size-independent harvest selection on fish body shape in two recreationally exploited marine species. Ecol. Evol., 4, 2154–2164.

Andersen, K.H., Berge, T., Gonçalves, R.J., Hartvig, M., Heuschele, J., Hylander, S., et al. (2016). Characteristic Sizes of Life in the Oceans, from Bacteria to Whales. Annu. Rev. Mar. Sci., 8, 217–241.

Berg, H.C. (1993). Random walks in biology. Princeton University Press.

Cermeño, P. & Figueiras, F.G. (2008). Species richness and cell-size distribution: size structure of phytoplankton communities. Mar. Ecol. Prog. Ser., 357, 79–85.

Charalampous, E., Matthiessen, B. & Sommer, U. (2018). Light effects on phytoplankton morphometric traits influence nutrient utilization ability. J. Plankton Res., 40, 568–579.

Dao, M.H. (2013). Reassessment of the cell surface area limitation to nutrient uptake in phytoplankton. Mar. Ecol. Prog. Ser., 489, 87–92.

Darwin, C. (1859). On the Origin of Species.

Durante, G., Basset, A., Stanca, E. & Roselli, L. (2019). Allometric scaling and morphological variation in sinking rate of phytoplankton. J. Phycol., 55, 1386–1393.

Edgington, H.A. & Taylor, D.R. (2019). Ecological contributions to body shape evolution in salamanders of the genus Eurycea (Plethodontidae). PLOS ONE, 14, e0216754.

Edwards, K.F., Thomas, M.K., Klausmeier, C.A. & Litchman, E. (2012). Allometric scaling and taxonomic variation in nutrient utilization traits and maximum growth rate of phytoplankton. Limnol Ocean., 57, 554–566.

Feldman, A., Sabath, N., Pyron, R.A., Mayrose, I. & Meiri, S. (2016). Body sizes and diversification rates of lizards, snakes, amphisbaenians and the tuatara. Glob. Ecol. Biogeogr., 25, 187–197.

Forsman, A. & Shine, R. (1995). Parallel Geographic Variation in Body Shape and Reproductive Life History within the Australian Scincid Lizard Lampropholis delicata. Funct. Ecol., 9, 818–828.

Gallet, R., Violle, C., Fromin, N., Jabbour-Zahab, R., Enquist, B.J. & Lenormand, T. (2017). The evolution of bacterial cell size: the internal diffusion-constraint hypothesis. ISME J., 11, 1559–1568.

Grover, J.P. (1989). Influence of Cell Shape and Size on Algal Competitive Ability. J. Phycol., 25, 402– 405.

Guiry, M.D. & Guiry, G.M. (2018). AlgaeBase. World-wide electronic publication, National University of Ireland, Galway. http://www.algaebase.org. AlgaeBase.

Helcom, H.C. (1988). Guidelines for the Baltic Monitoring Programme for the third stage. Balt. Sea Env. Proc D, 27, 1–60.

Hillebrand, H., Blasius, B., Borer, E.T., Chase, J.M., Downing, J.A., Eriksson, B.K., et al. (2018). Biodiversity change is uncoupled from species richness trends: Consequences for conservation and monitoring. J. Appl. Ecol., 55, 169–184.

Hillebrand, H., Dürselen, C.-D., Kirschtel, D., Pollingher, U. & Zohary, T. (1999). Biovolume calculation for pelagic and benthic microalgae. J. Phycol., 35, 403–424.

Hirst, A.G., Glazier, D.S. & Atkinson, D. (2014). Body shape shifting during growth permits tests that distinguish between competing geometric theories of metabolic scaling. Ecol. Lett., 17, 1274–1281.

Husemann, M., Tobler, M., McCauley, C., Ding, B. & Danley, P.D. (2017). Body shape differences in a pair of closely related Malawi cichlids and their hybrids: Effects of genetic variation, phenotypic plasticity, and transgressive segregation. Ecol. Evol., 7, 4336–4346.

ICES CEIM. (n.d.). Plankton cell sizes. Mar. Data. Available at: http://ices.dk/data/Documents/Forms/AllItems.aspx?RootFolder=%2fdata%2fDocuments%2fENV&FolderCTID=0x012000ACF6FBA45737584389AD23DD43BB914C. Last accessed.

Ignatiades, L. (2017). Size scaling patterns of species richness and carbon biomass for marine phytoplankton functional groups. Mar. Ecol., 38, e12454.

Karp-Boss, L. & Boss, E. (2016). The Elongated, the Squat and the Spherical: Selective Pressures for Phytoplankton Shape. In: Aquatic Microbial Ecology and Biogeochemistry: A Dual Perspective. Springer, pp. 25–34.

Kerimoglu, O., Straile, D. & Peeters, F. (2012). Role of phytoplankton cell size on the competition for nutrients and light in incompletely mixed systems. J. Theor. Biol., 300, 330–343.

Kiørboe, T. (2008). A mechanistic approach to plankton ecology. Princeton University Press.

Kruk, C. & Segura, A.M. (2012). The habitat template of phytoplankton morphology-based functional groups. Hydrobiologia, 698, 191–202.

Kumar, M., Mommer, M.S. & Sourjik, V. (2010). Mobility of cytoplasmic, membrane, and DNA-binding proteins in Escherichia coli. Biophys. J., 98, 552–559.

Lewis, W.M. (1976). Surface/volume ratio: implications for phytoplankton morphology. Science, 192, 885–887.

Li, Y., Ren, B., Ding, L., Shen, Q., Peng, S. & Guo, S. (2013). Does chloroplast size influence photosynthetic nitrogen use efficiency? PloS One, 8, e62036.

LifeWatch ERIC. (n.d.). LifeWatch Italy Data Portal. Available at: https://dataportal.lifewatchitaly.eu/data. Last accessed.

Litchman, E., Klausmeier, C.A. & Yoshiyama, K. (2009). Contrasting size evolution in marine and freshwater diatoms. Proc. Natl. Acad. Sci., 106, 2665–2670.

Malviya, S., Scalco, E., Audic, S., Vincent, F., Veluchamy, A., Poulain, J., et al. (2016). Insights into global diatom distribution and diversity in the world’s ocean. Proc. Natl. Acad. Sci., 113, E1516–E1525.

Marañón, E. (2015). Cell size as a key determinant of phytoplankton metabolism and community structure. Annu. Rev. Mar. Sci., 7, 241–264.

Martin, R.A. (2017). Body size in (mostly) mammals: mass, speciation rates and the translation of gamma to alpha diversity on evolutionary timescales. Hist. Biol., 29, 576–593.

Martin-Jézéquel, V., Hildebrand, M. & Brzezinski, M.A. (2000). Silicon metabolism in diatoms: implications for growth. J. Phycol., 36, 821–840.

May, R.M. (1986). The Search for Patterns in the Balance of Nature: Advances and Retreats. Ecology, 67, 1115–1126.

Milo, R. & Phillips, R. (2020). Cell Biology by then Numbers. Available at: book.bionumbers.org/what-are-the-time-scales-for-diffusion-in-cells/. Last accessed.

Mittler, U., Blasius, B., Gaedke, U. & Ryabov, A.B. (2019). Length-volume relationship of lake phytoplankton: Length-volume relationship of lake phytoplankton. Limnol. Oceanogr. Methods, 17, 58–68.

Monteiro, F.M., Bach, L.T., Brownlee, C., Bown, P., Rickaby, R.E.M., Poulton, A.J., et al. (2016). Why marine phytoplankton calcify. Sci. Adv., 2, e1501822.

Naselli-Flores, L. & Barone, R. (2007). Pluriannual morphological variability of phytoplankton in a highly productive Mediterranean reservoir (Lake Arancio, Southwestern Sicily). Hydrobiologia, 578, 87–95.

Naselli-Flores, L. & Barone, R. (2011). Invited Review -Fight on Plankton! or, Phytoplankton Shape and Size as Adaptive Tools to Get Ahead in the Struggle for Life. Cryptogam. Algol., 32, 157– 204.

Naselli-Flores, L., Padisák, J. & Albay, M. (2007). Shape and size in phytoplankton ecology: do they matter? Hydrobiologia, 578, 157–161.

Naselli-Flores, L., Zohary, T. & Padisák, J. (2020). Life in suspension and its impact on phytoplankton morphology: an homage to Colin S. Reynolds. Hydrobiologia.

Nelson, D.M., Tréguer, P., Brzezinski, M.A., Leynaert, A. & Quéguiner, B. (1995). Production and dissolution of biogenic silica in the ocean: revised global estimates, comparison with regional data and relationship to biogenic sedimentation. Glob. Biogeochem. Cycles, 9, 359–372.

Niklas, K.J. (2000). The evolution of plant body plans—a biomechanical perspective. Ann. Bot., 85, 411–438.

O’Farrell, I., de Tezanos Pinto, P. & Izaguirre, I. (2007). Phytoplankton morphological response to the underwater light conditions in a vegetated wetland. Hydrobiologia, 578, 65–77.

Olenina, I., Hajdu, S., Edler, L., Andersson, A., Wasmund, N., Busch, S., et al. (2006). Biovolumes and size-classes of phytoplankton in the Baltic Sea. HELCOM Balt.Sea Environ. Proc., 106.

Padisák, J., Soróczki-Pintér, É. & Rezner, Z. (2003). Sinking properties of some phytoplankton shapes and the relation of form resistance to morphological diversity of plankton—an experimental study. In: Aquatic biodiversity. Springer, pp. 243–257.

Pančić, M. & Kiørboe, T. (2018). Phytoplankton defence mechanisms: traits and trade-offs: Defensive traits and trade-offs. Biol. Rev., 93, 1269–1303.

Pawar, S., Dell, A.I. & Van M. Savage. (2012). Dimensionality of consumer search space drives trophic interaction strengths. Nature, 486, 485–489.

Pomati, F., Jokela, J., Simona, M., Veronesi, M. & Ibelings, B.W. (2011). An automated platform for phytoplankton ecology and aquatic ecosystem monitoring. Environ. Sci. Technol., 45, 9658– 9665.

Porter, W.P. & Kearney, M. (2009). Size, shape, and the thermal niche of endotherms. Proc. Natl. Acad. Sci., 106, 19666–19672.

Raven, J.A. (1998). The twelfth Tansley Lecture. Small is beautiful: the picophytoplankton. Funct. Ecol., 12, 503–513.

Reynolds, C.S. (1988). Functional morphology and the adaptive strategies of freshwater phytoplankton. In: Growth and reproductive strategies of freshwater phytoplankton. Cambridge University Press, pp. 388–433.

Reynolds, C.S. (2006). The Ecology of Phytoplankton. Ecology, Biodiversity and Conservation. Cambridge University Press, Cambridge.

Roselli, L., Litchman, E., Elena, S., Cozzoli, F. & Basset, A. (2017). Individual trait variation in phytoplankton communities across multiple spatial scales. J. Plankton Res., 39, 577–588.

Ryabov, A.B. & Blasius, B. (2011). A graphical theory of competition on spatial resource gradients. Ecol Lett, 14, 220–228.

Salmaso, N. & Padisák, J. (2007). Morpho-functional groups and phytoplankton development in two deep lakes (Lake Garda, Italy and Lake Stechlin, Germany). Hydrobiologia, 578, 97–112.

Siemann, E., Tilman, D. & Haarstad, J. (1996). Insect species diversity, abundance and body size relationships. Nature, 380, 704–706.

Stanca, E., Cellamare, M. & Basset, A. (2013). Geometric shape as a trait to study phytoplankton distributions in aquatic ecosystems. Hydrobiologia, 701, 99–116.

Sunda, W.G. & Hardison, D.R. (2010). Evolutionary tradeoffs among nutrient acquisition, cell size, and grazing defense in marine phytoplankton promote ecosystem stability. Mar. Ecol. Prog. Ser., 401, 63–76.

Tambi, H., Flaten, G.A.F., Egge, J.K., Bødtker, G., Jacobsen, A. & Thingstad, T.F. (2009). Relationship between phosphate affinities and cell size and shape in various bacteria and phytoplankton. Aquat. Microb. Ecol., 57, 311–320.

Utermöhl, H. (1958). Zur vervollkommnung der quantitativen phytoplankton-methodik: Mit 1 Tabelle und 15 abbildungen im Text und auf 1 Tafel. Int. Ver. Für Theor. Angew. Limnol. Mitteilungen, 9, 1–38.

Vadrucci, M.R., Cabrini, M. & Basset, A. (2007). Biovolume determination of phytoplankton guilds in transitional water ecosystems of Mediterranean Ecoregion. Transitional Waters Bull., 1, 83– 102.

Weeks, B.C., Willard, D.E., Zimova, M., Ellis, A.A., Witynski, M.L., Hennen, M., et al. (2020). Shared morphological consequences of global warming in North American migratory birds. Ecol. Lett., 23, 316–325.

Weithoff, G. & Gaedke, U. (2017). Mean functional traits of lake phytoplankton reflect seasonal and inter-annual changes in nutrients, climate and herbivory. J. Plankton Res., 509–517.

Willén, T. (1962). Studies on the phytoplankton of some lakes connected with or recently isolated from the Baltic. Oikos, 13, 169–199.

Wirtz, K.W. (2011). Non-uniform scaling in phytoplankton growth rate due to intracellular light and CO2 decline. J. Plankton Res., 33, 1325–1341.

Zohary, T., Padisák, J. & Naselli-Flores, L. (2010). Phytoplankton in the physical environment: beyond nutrients, at the end, there is some light. Hydrobiologia, 639, 261–269.

