## Supplementary material for "Shape matters: the relationship between cell geometry and diversity in phytoplankton"

### Initialize

*restart*;

Radius
$R := 'R';$

$$R := R \quad (1.1)$$

Semi axis
$A := 'A'; B := 'B'; C := 'C';$

$$A := A$$
$$B := B$$
$$C := C \quad (1.2)$$

Height
$H := 'H';$

$$H := H \quad (1.3)$$

### Elementary shapes, 2D

$Circle\_A := \mathrm{pi} \cdot R^2;$

$$Circle\_A := \pi R^2 \quad (2.1)$$

$Circle\_P := 2 \cdot \mathrm{pi} \cdot R;$

$$Circle\_P := 2 \pi R \quad (2.2)$$

### Elementary shapes, 3D

$Sphere\_A := 4 \cdot \mathrm{pi} \cdot R^2;$

$$Sphere\_A := 4 \pi R^2 \quad (3.1)$$

$Sphere\_V := \frac{4}{3} \cdot \mathrm{pi} \cdot R^3$

$$Sphere\_V := \frac{4 \pi R^3}{3} \quad (3.2)$$

$Cone\_V := \frac{1}{3} \cdot \mathrm{pi} \cdot R^2 \cdot H;$

$$Cone\_V := \frac{\pi R^2 H}{3} \quad (3.3)$$

$Cone\_Atop := \mathrm{pi} \cdot R^2;$

$$Cone\_Atop := \pi R^2 \quad (3.4)$$

$Cone\_ASide := \mathrm{pi} \cdot R \cdot \mathrm{sqrt}(R^2 + H^2)$

$$Cone\_ASide := \pi R \sqrt{H^2 + R^2} \tag{3.5}$$

$Cone\_A := Cone\_Atop + Cone\_ASide;$

$$Cone\_A := \pi R^2 + \pi R \sqrt{H^2 + R^2} \tag{3.6}$$

$Cylinder\_V := \mathrm{pi} \cdot R^2 \cdot H;$

$$Cylinder\_V := \pi R^2 H \tag{3.7}$$

$Cylinder\_Side\_A := 2 \cdot \mathrm{pi} \cdot R \cdot H;$

$$Cylinder\_Side\_A := 2 \pi R H \tag{3.8}$$

$Cylinder\_BaseTop\_A := 2 \cdot \mathrm{pi} \cdot R^2;$

$$Cylinder\_BaseTop\_A := 2 \pi R^2 \tag{3.9}$$

$Cylinder\_A := Cylinder\_Side\_A + Cylinder\_BaseTop\_A;$

$$Cylinder\_A := 2 \pi R H + 2 \pi R^2 \tag{3.10}$$

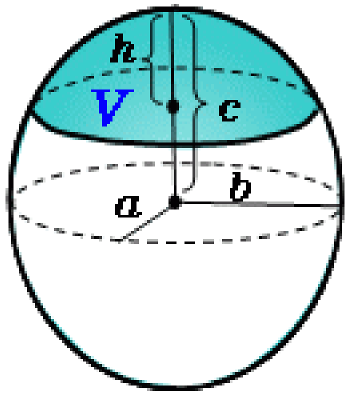


Ellipsoidal segment (http://keisan.casio.com/exec/system/1311572253)

### Subs R=d/2

$R := \frac{d}{2};$

$$R := \frac{d}{2} \tag{4.1}$$

$A := \frac{a}{2}; B := \frac{b}{2}; C := \frac{c}{2};$

$$A := \frac{a}{2}$$

$$B := \frac{b}{2}$$

$$C := \frac{c}{2} \tag{4.2}$$

### Ellipse

$Ellipse\_A := \text{pi}\cdot A\cdot B;$

$$Ellipse\_A := \frac{\pi a b}{4} \tag{5.1}$$

$Q := \frac{(A-B)^2}{(A+B)^2};$

$$Q := \frac{\left(\frac{a}{2}-\frac{b}{2}\right)^2}{\left(\frac{a}{2}+\frac{b}{2}\right)^2} \tag{5.2}$$

$Q := normal(Q);$

$$Q := \frac{(a-b)^2}{(a+b)^2} \tag{5.3}$$

$Ellipse\_P := \text{pi}\cdot(A+B)\cdot\left(1+\frac{1}{4}\cdot Q\right);$

$$Ellipse\_P := \pi\left(\frac{a}{2}+\frac{b}{2}\right)\left(1+\frac{(a-b)^2}{4\,(a+b)^2}\right) \tag{5.4}$$

### Simple shapes, 3D (as a function of d, a, b, c)

$Sphere\_A$

$$\pi d^2 \tag{6.1}$$

$Sphere\_V$

$$\frac{\pi d^3}{6} \tag{6.2}$$

Note that for ellipsoid I already use b and c instead of the semi axes A and B

$Ellipsoid\_V := \text{pi}/6\cdot b\cdot c\cdot H;$

$$Ellipsoid\_V := \frac{\pi b c H}{6} \tag{6.3}$$

This formula works only for prolate ellipsoid (2H>b+c)

$Ellipsoid\_A :=\ \text{pi}/4\cdot(b+c)\cdot(\ (b+c)/2+2\cdot H\hat{}2/\text{sqrt}(4\cdot H\hat{}2-(b+c)\hat{}2)$
$\cdot\ \text{arcsin}(\text{sqrt}(4\cdot H\hat{}2-(b+c)\hat{}2)/(2\cdot H)\ )\ );$

$$Ellipsoid\_A := \frac{\pi\,(b+c)\left(\frac{b}{2}+\frac{c}{2}+\frac{2 H^2 \arcsin\left(\frac{\sqrt{4 H^2-(b+c)^2}}{2 H}\right)}{\sqrt{4 H^2-(b+c)^2}}\right)}{4} \tag{6.4}$$

Knud Thomsen’s Formula is better, because works for any allipsoid

$$Ellipsoid\_KTA := 4\cdot \mathrm{pi}\cdot\left(\frac{A^pB^p + A^p\cdot\left(\frac{H}{2}\right)^p + B^p\cdot\left(\frac{H}{2}\right)^p}{3}\right)^{\frac{1}{p}}$$

$$Ellipsoid\_KTA := 4\,\pi\left(\frac{\left(\frac{a}{2}\right)^p\left(\frac{b}{2}\right)^p}{3} + \frac{\left(\frac{a}{2}\right)^p\left(\frac{H}{2}\right)^p}{3} + \frac{\left(\frac{b}{2}\right)^p\left(\frac{H}{2}\right)^p}{3}\right)^{\frac{1}{p}} \tag{6.5}$$

$$RotEllipsoid\_A := 2\cdot \mathrm{pi}\cdot A\left(A + \frac{H^2}{\mathrm{sqrt}(H^2 - C^2)}\cdot \arcsin\left(\frac{\mathrm{sqrt}(H^2 - A^2)}{H}\right)\right);$$

$$RotEllipsoid\_A := \pi\, a\left(\frac{a}{2} + \frac{2\,H^2 \arcsin\left(\frac{\sqrt{4\,H^2 - a^2}}{2\,H}\right)}{\sqrt{4\,H^2 - c^2}}\right) \tag{6.6}$$

$$Ellipsoid\_KTA := 4\cdot \mathrm{pi}\cdot\left(\frac{A^pB^p + A^p\cdot C^p + B^p\cdot C^p}{3}\right)^{\frac{1}{p}}$$

$$Ellipsoid\_KTA := 4\,\pi\left(\frac{\left(\frac{a}{2}\right)^p\left(\frac{b}{2}\right)^p}{3} + \frac{\left(\frac{a}{2}\right)^p\left(\frac{c}{2}\right)^p}{3} + \frac{\left(\frac{b}{2}\right)^p\left(\frac{c}{2}\right)^p}{3}\right)^{\frac{1}{p}} \tag{6.7}$$

$Spheroid\_V := subs(b = d, c = d, Ellipsoid\_V);$

$$Spheroid\_V := \frac{\pi\, d^2\, H}{6} \tag{6.8}$$

$Spheroid\_A := (expand(simplify(subs(b = d, c = d, Ellipsoid\_A))));$

$$Spheroid\_A := \frac{\pi\, d\, H^2 \arcsin\left(\frac{\sqrt{H^2 - d^2}}{H}\right)}{2\sqrt{H^2 - d^2}} + \frac{\pi\, d^2}{2} \tag{6.9}$$

$Cylinder\_V;$

$$\frac{\pi\, d^2\, H}{4} \tag{6.10}$$

$Cylinder\_A;$

$$H\, d\, \pi + \frac{1}{2}\,\pi\, d^2 \tag{6.11}$$

$Cone\_V$

$$\frac{\pi\, d^2\, H}{12} \tag{6.12}$$

$Cone\_Atop$

$$\frac{\pi\, d^2}{4} \tag{6.13}$$

$simplify(Cone\_ASide)$

$$\frac{\pi d \sqrt{4 H^2 + d^2}}{4} \quad (6.14)$$

(*simplify*(*Cone_A*));

$$\frac{\pi d \left(d + \sqrt{4 H^2 + d^2}\right)}{4} \quad (6.15)$$

(*EllipticPrism_A*);

$$EllipticPrism\_A \quad (6.16)$$

*simplify*(*EllipticPrism_V*)

$$EllipticPrism\_V \quad (6.17)$$

### Complex shapes

#### 1. Sphere

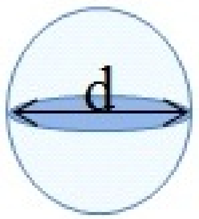


*A_1* := *Sphere_A*

$$A\_1 := \pi d^2 \quad (7.1.1)$$

*V_1* := *Sphere_V*

$$V\_1 := \frac{\pi d^3}{6} \quad (7.1.2)$$

#### 2. Prolate spheroid

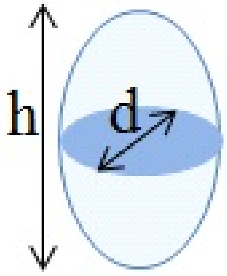


*A_2* := *subs*(*c* = *d*, *b* = *d*, *H* = *h*, *Ellipsoid_A*)

$$A\_2 := \frac{\pi\, d \left( d + \frac{2\, h^2 \arcsin\left( \frac{\sqrt{-4\, d^2 + 4\, h^2}}{2\, h} \right)}{\sqrt{-4\, d^2 + 4\, h^2}} \right)}{2} \quad \textbf{(7.2.1)}$$

$V\_2 := subs(c = d, b = d, H = h, Ellipsoid\_V)$

$$V\_2 := \frac{\pi\, d^2\, h}{6} \quad \textbf{(7.2.2)}$$

#### 3.Cylinder

$A\_3 := subs(H = h, Cylinder\_A);$

$$A\_3 := h\, d\, \pi + \frac{1}{2}\, \pi\, d^2 \quad \textbf{(7.3.1)}$$

$V\_3 := subs(H = h, Cylinder\_V)$

$$V\_3 := \frac{\pi\, d^2\, h}{4} \quad \textbf{(7.3.2)}$$

#### 4. Ellipsoid

$A\_4 := subs(c = c, b = b, H = h, Ellipsoid\_A)$

$$A\_4 := \frac{\pi\, (b + c) \left( \frac{b}{2} + \frac{c}{2} + \frac{2\, h^2 \arcsin\left( \frac{\sqrt{4\, h^2 - (b + c)^2}}{2\, h} \right)}{\sqrt{4\, h^2 - (b + c)^2}} \right)}{4} \quad \textbf{(7.4.1)}$$

$V\_4 := subs(c = c, b = b, H = h, Ellipsoid\_V)$

$$V\_4 := \frac{\pi\, b\, c\, h}{6} \quad \textbf{(7.4.2)}$$

#### 5. Cone

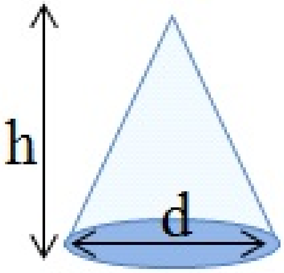


$A\_5 := expand(simplify(subs(H = h, Cone\_A)));$

$$A\_5 := \frac{\pi d^2}{4} + \frac{\pi d \sqrt{d^2 + 4 h^2}}{4} \tag{7.5.1}$$

$V\_5 := subs(H = h, Cone\_V);$

$$V\_5 := \frac{\pi d^2 h}{12} \tag{7.5.2}$$

### 6. Truncated cone

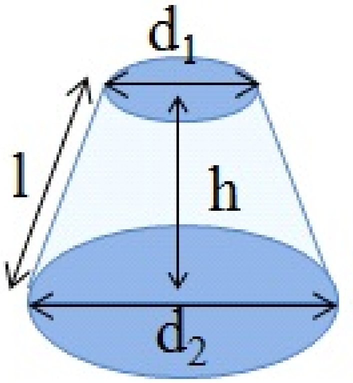


Formulas for the area and volume from http://keisan.casio.com/exec/system/1223372110

$Cone\_Trunc\_A\_Side := \mathrm{pi} \cdot (R1 + R2) \cdot \mathrm{sqrt}((R2 - R1)^2 + h^2);$

$$Cone\_Trunc\_A\_Side := \pi (R1 + R2) \sqrt{(R2 - R1)^2 + h^2} \tag{7.6.1}$$

$Cone\_Trunc\_A\_Side := subs\left(R1 = \frac{d1}{2}, R2 = \frac{d2}{2}, Cone\_Trunc\_A\_Side\right);$

$$Cone\_Trunc\_A\_Side := \pi \left(\frac{d1}{2} + \frac{d2}{2}\right) \sqrt{\left(\frac{d2}{2} - \frac{d1}{2}\right)^2 + h^2} \tag{7.6.2}$$

$Cone\_Trunc\_A\_Base := subs(d = d1, Circle\_A) + subs(d = d2, Circle\_A);$

$$Cone\_Trunc\_A\_Base := \frac{1}{4} \pi d1^2 + \frac{1}{4} \pi d2^2 \tag{7.6.3}$$

$Cone\_Trunc\_A := Cone\_Trunc\_A\_Side + Cone\_Trunc\_A\_Base;$

$$Cone\_Trunc\_A := \pi \left(\frac{d1}{2} + \frac{d2}{2}\right) \sqrt{\left(\frac{d2}{2} - \frac{d1}{2}\right)^2 + h^2} + \frac{\pi d1^2}{4} + \frac{\pi d2^2}{4} \tag{7.6.4}$$

$Cone\_Trunc\_V := \frac{\mathrm{pi} \cdot h \cdot (R1^2 + R1 \cdot R2 + R2^2)}{3};$

$$Cone\_Trunc\_V := \frac{\pi h (R1^2 + R2 R1 + R2^2)}{3} \tag{7.6.5}$$

$Cone\_Trunc\_V := simplify\left(subs\left(R1 = \frac{d1}{2}, R2 = \frac{d2}{2}, Cone\_Trunc\_V\right)\right)$

$$Cone\_Trunc\_V := \frac{\pi h (d1^2 + d2 d1 + d2^2)}{12} \tag{7.6.6}$$

$A\_8 := Cone\_Trunc\_A;$

$$A\_8 := \pi \left(\frac{d1}{2} + \frac{d2}{2}\right) \sqrt{\left(\frac{d2}{2} - \frac{d1}{2}\right)^2 + h^2} + \frac{\pi\, d1^2}{4} + \frac{\pi\, d2^2}{4} \qquad (7.6.7)$$

$V\_8 := Cone\_Trunc\_V;$

$$V\_8 := \frac{\pi\, h\, (d1^2 + d2\, d1 + d2^2)}{12} \qquad (7.6.8)$$

#### 7 Parallelepiped

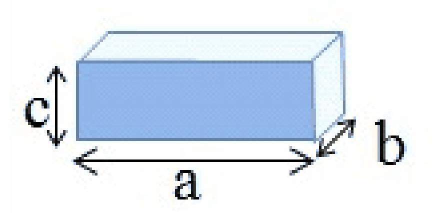


$A\_7 := 2 * a * b + 2 * b * c + 2 * a * c$

$$A\_7 := 2\, a\, b + 2\, a\, c + 2\, b\, c \qquad (7.7.1)$$

$V\_7 := a \cdot b \cdot c;$

$$V\_7 := a\, b\, c \qquad (7.7.2)$$

#### 8. Prism on elliptic base

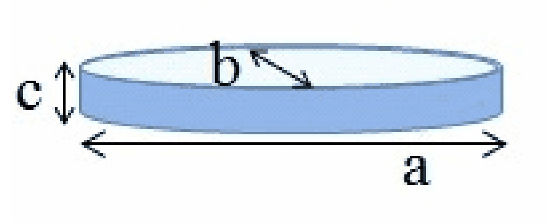


$Ellipse\_P$

$$\pi \left(\frac{a}{2} + \frac{b}{2}\right) \left(1 + \frac{(a-b)^2}{4\, (a+b)^2}\right) \qquad (7.8.1)$$

$EllipticPrism\_A := 2 \cdot Ellipse\_A + Ellipse\_P \cdot c;$

$$EllipticPrism\_A := c\, \pi \left(\frac{a}{2} + \frac{b}{2}\right) \left(1 + \frac{(a-b)^2}{4\, (a+b)^2}\right) + \frac{\pi\, a\, b}{2} \qquad (7.8.2)$$

$EllipticPrism\_V := Ellipse\_A \cdot c;$

$$EllipticPrism\_V := \frac{\pi\, a\, b\, c}{4} \qquad (7.8.3)$$

$evalf(subs(\mathrm{pi} = \mathrm{Pi}, a = 98.548, b = 7.06297781581347, c = 3.28482, EllipticPrism\_A))$

$$1740.496737 \qquad (7.8.4)$$

#### 9. Prism on parallelogram base

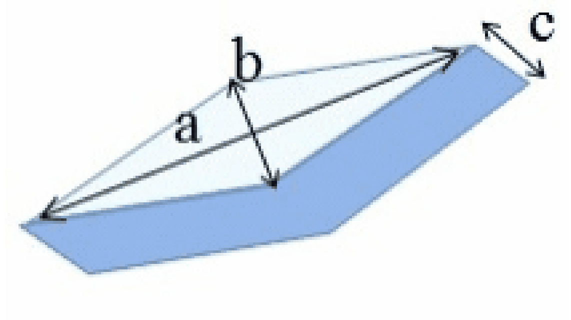


$A\_9 := a*b + 2*\mathrm{sqrt}(a\char`^2 + b\char`^2)*c$

$$A\_9 := a\,b + 2\sqrt{a^2 + b^2}\,c \quad \textbf{(7.9.1)}$$

$V\_9 := 1/2*a*b*c$

$$V\_9 := \frac{a\,b\,c}{2} \quad \textbf{(7.9.2)}$$

#### 10. Cube

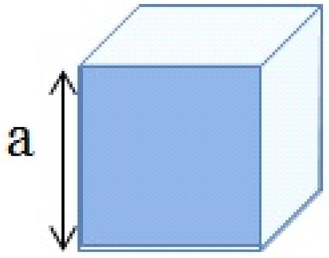


$A\_10 := 6\cdot a^2$

$$A\_10 := 6\,a^2 \quad \textbf{(7.10.1)}$$

$V\_10 := a^3$

$$V\_10 := a^3 \quad \textbf{(7.10.2)}$$

#### 11 Prism on triangle base 1

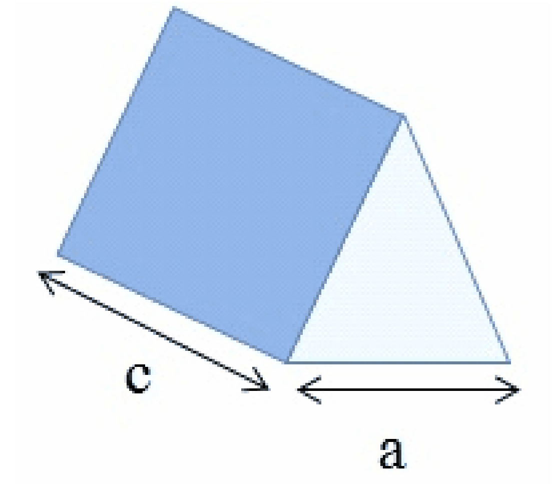


A_11 := 3 a·c + sqrt(3) / 2 * a^2

$$A\_11 := 3\,a\,c + \frac{\sqrt{3}\,a^2}{2} \tag{7.11.1}$$

V_11 := sqrt(3) / 4 · a^2 * c

$$V\_11 := \frac{\sqrt{3}\,a^2\,c}{4} \tag{7.11.2}$$

#### 12 Half prism on elliptic base

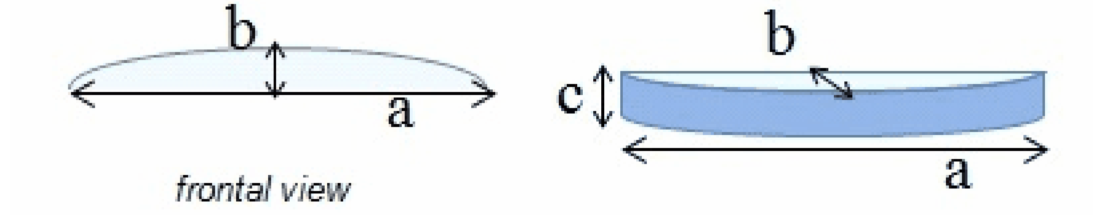


Area = 1/2 EllipticCylinder Side + Rectrangular_Side + 2 * 1/2 * EllipseArea

$$A\_12 := \frac{1}{2} \cdot subs(a = a, b = 2 \cdot b, Ellipse\_P) \cdot c + a \cdot c + 2 \cdot \frac{1}{2} \cdot subs(a = a, b = 2 \cdot b, Ellipse\_A)$$

$$A\_12 := \frac{\pi \left(\frac{a}{2} + b\right) \left(1 + \frac{(a - 2\,b)^2}{4\,(a + 2\,b)^2}\right) c}{2} + a\,c + \frac{\pi\,a\,b}{2} \tag{7.12.1}$$

$$A\_12\_simpl := simplify\left(\frac{\text{pi}\left(\frac{a}{2} + b\right)\ c}{2} + a\,c + \frac{\text{pi}\,a\,b}{2}\right)$$

$$A\_12\_simpl := \frac{((2\,b + c)\,a + 2\,b\,c)\,\pi}{4} + a\,c \tag{7.12.2}$$

$$V\_12 := \frac{1}{2} \cdot subs(a = a, b = 2 \cdot b, Ellipse\_A) \cdot c;$$

$$V\_12 := \frac{\pi\,a\,b\,c}{4} \tag{7.12.3}$$

#### 14. Cone + Cone (AtSh has the same)

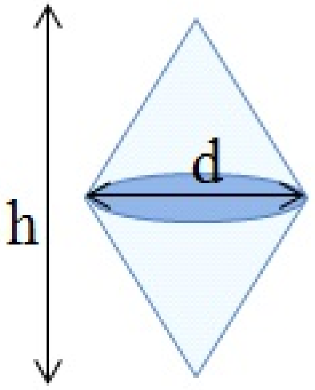


$A\_14 := simplify\left(2 \cdot subs\left(H = \frac{h}{2}, Cone\_ASide\right)\right)$

$$A\_14 := \frac{\pi\, d \sqrt{d^2 + h^2}}{2} \quad (7.13.1)$$

$V\_14 := simplify\left(2 \cdot subs\left(H = \frac{h}{2}, Cone\_V\right)\right)$

$$V\_14 := \frac{\pi\, d^2\, h}{12} \quad (7.13.2)$$

#### 15. Truncated cone + Truncated cone

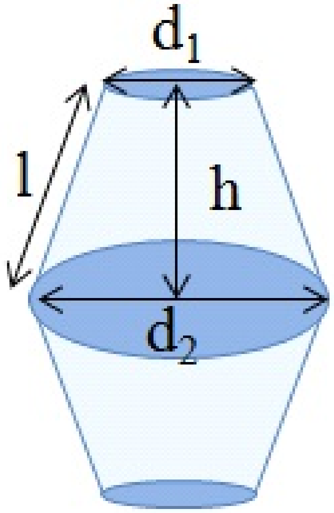


I didn't check it yet. There is only one record

#### 16. Prolate spheroid + 2 Cylinders

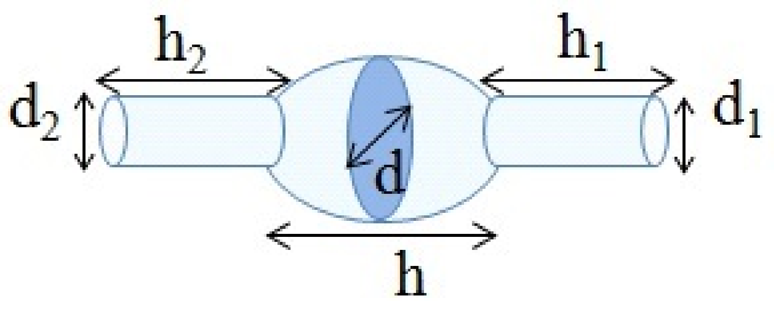


$$d_1 = d_2$$

Area = CylinderSide1 + CylinderSide2 + EllipsoidArea. (Cylinder Top is not included, because it is approximately the area closed by the cylinder bottoms on the ellipsoids)

$A\_16 = subs(H = h1, d = d1, Cylinder\_Side\_A) + subs(H = h2, d = d2, Cylinder\_Side\_A) + subs(d = d, H = h, Spheroid\_A)$

$$A\_16 = \pi\, d1\, h1 + \pi\, d2\, h2 + \frac{\pi\, d\, h^2 \arcsin\left(\frac{\sqrt{-d^2 + h^2}}{h}\right)}{2\sqrt{-d^2 + h^2}} + \frac{\pi\, d^2}{2} \qquad \textbf{(7.15.1)}$$

$V\_16 = subs(H = h1, d = d1, Cylinder\_V) + subs(H = h2, d = d2, Cylinder\_V) + subs(d = d, H = h, Spheroid\_V)$

$$V\_16 = \frac{1}{4}\pi\, d1^2\, h1 + \frac{1}{4}\pi\, d2^2\, h2 + \frac{1}{6}\pi\, d^2\, h \qquad \textbf{(7.15.2)}$$

#### 17. Cylinder + 2 Cones

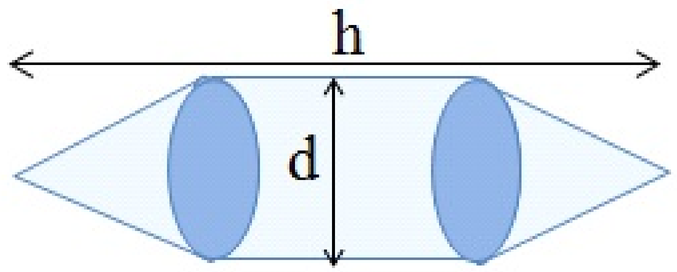


The formula assumes that h is the cylinder size and the cones are equilateral so their height is

$h1 = \frac{\mathrm{sqrt}(3)}{2} \cdot d$

$$h1 = \frac{\sqrt{3}\, d}{2} \qquad \textbf{(7.16.1)}$$

$A\_17b := 2 \cdot \frac{simplify\left(subs\left(H = \frac{\mathrm{sqrt}(3)}{2} \cdot d, Cone\_ASide\right)\right)}{\mathrm{csgn}(d)} + subs\left(H = h - 2 \cdot \frac{\mathrm{sqrt}(3)}{2} d,\right.$

$Cylinder\_Side\_A\Big)$

$$A\_17b := \pi d^2 + \pi d \left(h - \sqrt{3}\, d\right) \qquad \textbf{(7.16.2)}$$

$V\_17b := 2 \cdot simplify\left(subs\left(H = \frac{\mathrm{sqrt}(3)}{2} \cdot d, Cone\_V\right)\right) + subs\Big(H = h - 2 \cdot \frac{\mathrm{sqrt}(3)}{2} d,$
$Cylinder\_V\Big)$

$$V\_17b := \frac{\pi d^3 \sqrt{3}}{12} + \frac{\pi d^2 \left(h - \sqrt{3}\, d\right)}{4} \qquad \textbf{(7.16.3)}$$

Here I assume that the cone height h1=d/2 and the unit height is h  as suggested in Sun03

$A\_17 := 2 \cdot \frac{simplify\left(subs\left(H = \frac{1}{2} \cdot d, Cone\_ASide\right)\right)}{\mathrm{csgn}(d)} + subs(H = h - d, Cylinder\_Side\_A)$

$$A\_17 := \frac{\pi d^2 \sqrt{2}}{2} + \pi d\,(h - d) \qquad \textbf{(7.16.4)}$$

$V\_17 := \left(expand\left(2 \cdot simplify\left(subs\left(H = \frac{1}{2} \cdot d, Cone\_V\right)\right) + subs(H = h - d, Cylinder\_V)\right)\right)$

$$V\_17 := -\frac{1}{6}\pi d^3 + \frac{1}{4}\pi d^2 h \qquad \textbf{(7.16.5)}$$

For the same paramters as in Hi99
$A\_17Hi99 := 2 \cdot simplify(subs(H = z, Cone\_ASide)) + subs(H = h, Cylinder\_Side\_A)$

$$A\_17Hi99 := \frac{\pi d \sqrt{d^2 + 4 z^2}}{2} + h d \pi \qquad \textbf{(7.16.6)}$$

$V\_17Hi99 := expand(simplify(2 \cdot simplify(subs(H = z, Cone\_V)) + subs(H = h, Cylinder\_V)))$

$$V\_17Hi99 := \frac{1}{6}\pi d^2 z + \frac{1}{4}\pi d^2 h \qquad \textbf{(7.16.7)}$$

#### 19. Cone + Half sphere

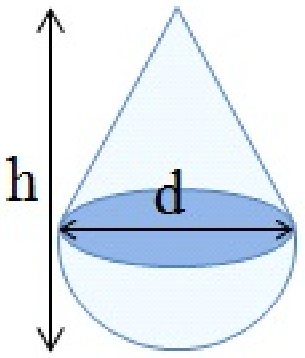


$A\_19 := factor\left(\left(subs\left(H = h - \frac{d}{2}, (Cone\_ASide)\right) + \frac{1}{2} \cdot Sphere\_A\right)\right)$

$$A\_19 := \frac{\pi\, d \left(\sqrt{2\, d^2 - 4\, h\, d + 4\, h^2} + 2\, d\right)}{4} \tag{7.17.1}$$

$V\_19 := simplify\left(simplify\left(subs\left(H = h - \frac{d}{2}, Cone\_V\right)\right) + \frac{1}{2} \cdot Sphere\_V\right)$

$$V\_19 := \frac{\pi\, d^2\, (d + 2\, h)}{24} \tag{7.17.2}$$

#### 20. Half ellipsoid + Cone (on elliptic base)

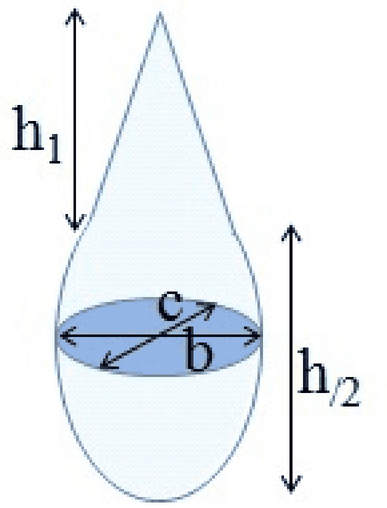


$Cone\_EllBase\_ASide := \frac{1}{2} \cdot \mathrm{pi} \cdot \left(B \cdot \mathrm{sqrt}\left(B^2 + h1^2\right) + C \cdot \mathrm{sqrt}\left(C^2 + h1^2\right)\right);$

$$Cone\_EllBase\_ASide := \frac{\pi \left(\frac{b \sqrt{b^2 + 4\, h1^2}}{4} + \frac{c \sqrt{c^2 + 4\, h1^2}}{4}\right)}{2} \tag{7.18.1}$$

Check if this formula gives a correct answer for the cone with circular base

$simplify(subs(b = d, c = d, h1 = H, Cone\_EllBase\_ASide) - Cone\_ASide);$

$$0 \tag{7.18.2}$$

$Cone\_EllBase\_V := \frac{1}{3} subs(a = c, b = b, Ellipse\_A) \cdot h1;$

$$Cone\_EllBase\_V := \frac{\pi\, c\, b\, h1}{12} \tag{7.18.3}$$

$A\_20 := Cone\_EllBase\_ASide + \frac{1}{2} subs(H = h, Ellipsoid\_A);$

$$A\_20 := \frac{\pi \left( \frac{b\sqrt{b^2+4\,h1^2}}{4} + \frac{c\sqrt{c^2+4\,h1^2}}{4} \right)}{2} + \frac{\pi\,(b+c)\left( \frac{b}{2} + \frac{c}{2} + \frac{2\,h^2 \arcsin\left( \frac{\sqrt{4\,h^2-(b+c)^2}}{2\,h} \right)}{\sqrt{4\,h^2-(b+c)^2}} \right)}{8} \qquad (7.18.4)$$

V_20:=

$Cone\_EllBase\_V + \frac{1}{2} subs(H=h, Ellipsoid\_V)$

$$\frac{1}{12}\,\pi\,c\,b\,h1 + \frac{1}{12}\,\pi\,b\,c\,h \qquad (7.18.5)$$

#### 21.Prism on elliptic base+ box

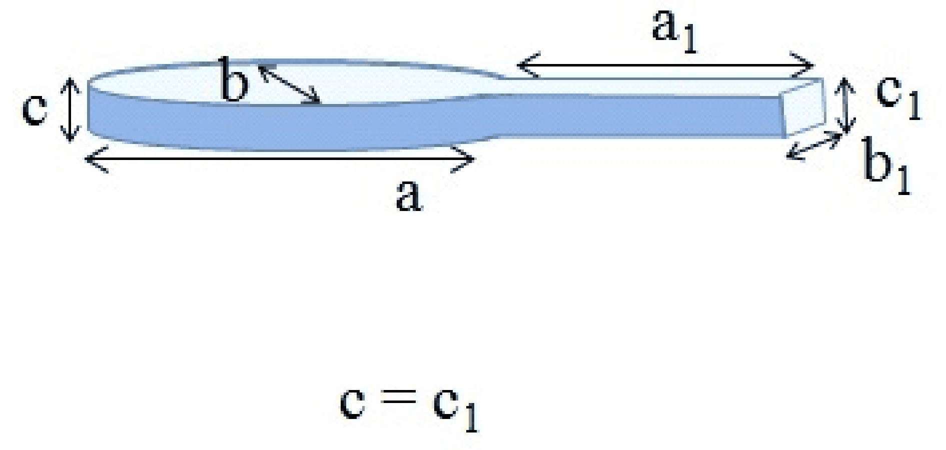


Shape perimeter. We should subtract b1 from the ellipse perimeter (web has an extra b1)
$P\_21 := Ellipse\_P - b1 + 2\cdot a1 + b1;$

$$P\_21 := \pi \left( \frac{a}{2} + \frac{b}{2} \right) \left( 1 + \frac{(a-b)^2}{4\,(a+b)^2} \right) + 2\,a1 \qquad (7.19.1)$$

shape volume, matches the Web

$V\_21 := EllipticPrism\_V + a1\cdot b1\cdot c;$

$$V\_21 := a1\,b1\,c + \frac{1}{4}\,\pi\,a\,b\,c \qquad (7.19.2)$$

Shape area (no)

$A\_21 := P\_21\cdot c + 2\cdot Ellipse\_A + 2\cdot a1\cdot b1;$

$$A\_21 := \left( \pi \left( \frac{a}{2} + \frac{b}{2} \right) \left( 1 + \frac{(a-b)^2}{4\,(a+b)^2} \right) + 2\,a1 \right) c + 2\,a1\,b1 + \frac{\pi\,a\,b}{2} \qquad (7.19.3)$$

#### 22. Cylinder + 2 Half spheres

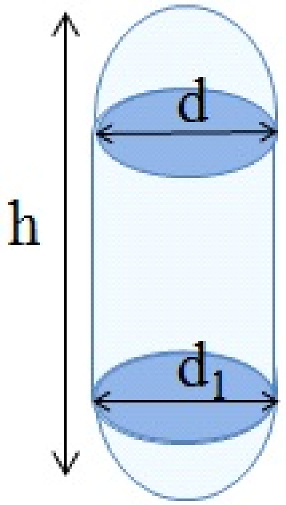


$d = d_1$

the figure is misleading, as it shows that h is the full height of the unit, but it is only the cylinder height

$A\_21 := 2 \cdot \frac{1}{2} \cdot Sphere\_A + subs(H = h, Cylinder\_Side\_A);$

$$A\_21 := \pi d^2 + h d \pi \quad \textbf{(7.20.1)}$$

$simplify(A\_21)$

$$\pi d (d + h) \quad \textbf{(7.20.2)}$$

$V\_21 := 2 \cdot \frac{1}{2} Sphere\_V + subs(H = h, Cylinder\_V)$

$$V\_21 := \frac{1}{6} \pi d^3 + \frac{1}{4} \pi d^2 h \quad \textbf{(7.20.3)}$$

$simplify(V\_21)$

$$\frac{\pi d^2 (2 d + 3 h)}{12} \quad \textbf{(7.20.4)}$$

#### 23. Ellipsoid+2cones+cylinder

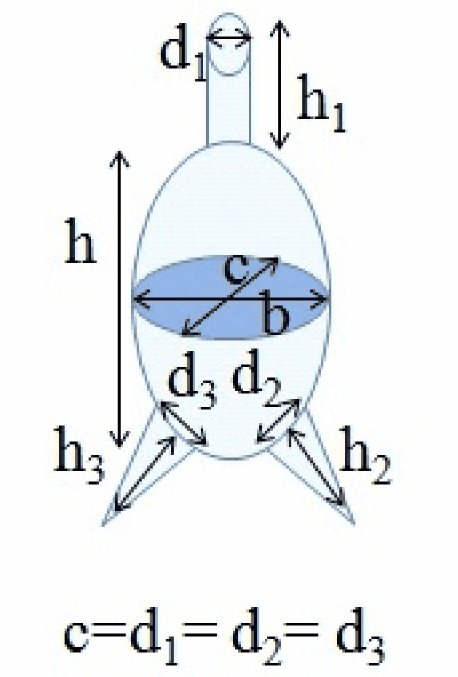


$A\_23 := subs(H=h, Ellipsoid\_A) - subs(d=d1, Circle\_A) - subs(d=d2, Circle\_A) - subs(d=d3, Circle\_A) + subs(d=d1, Circle\_A) + subs(H=h1, d=d1, Cylinder\_Side\_A) + subs(H=h2, d=d2, Cone\_ASide) + subs(H=h3, d=d3, Cone\_ASide);$

$$A\_23 := \frac{\pi (b+c)\left(\frac{b}{2}+\frac{c}{2}+\frac{2h^2 \arcsin\left(\frac{\sqrt{4h^2-(b+c)^2}}{2h}\right)}{\sqrt{4h^2-(b+c)^2}}\right)}{4} - \frac{\pi d2^2}{4} - \frac{\pi d3^2}{4} + \pi\, d1\, h1 + \frac{\pi\, d2 \sqrt{h2^2+\frac{d2^2}{4}}}{2} + \frac{\pi\, d3 \sqrt{h3^2+\frac{d3^2}{4}}}{2} \tag{7.21.1}$$

$V\_23 := subs(H=h, Ellipsoid\_V) + subs(H=h1, d=d1, Cylinder\_V) + subs(H=h2, d=d2, Cone\_V) + subs(H=h3, d=d3, Cone\_V)$

$$V\_23 := \frac{1}{6}\pi\, b\, c\, h + \frac{1}{4}\pi\, d1^2\, h1 + \frac{1}{12}\pi\, d2^2\, h2 + \frac{1}{12}\pi\, d3^2\, h3 \tag{7.21.2}$$

#### 24. Ellipsoid + Cone

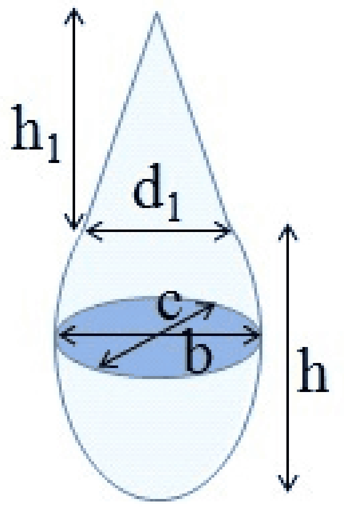


Area = EllipsoidArea - ConeBase + ConeSide

$A\_24 := subs(H=h, Ellipsoid\_A) - subs(d=d1, Circle\_A) + subs(H=h1, d=d1, Cone\_ASide)$

$$A\_24 := \frac{\pi (b+c) \left( \frac{b}{2} + \frac{c}{2} + \frac{2 h^2 \arcsin\left( \frac{\sqrt{4 h^2 - (b+c)^2}}{2 h} \right)}{\sqrt{4 h^2 - (b+c)^2}} \right)}{4} - \frac{\pi d1^2}{4} + \frac{\pi d1 \sqrt{h1^2 + \frac{d1^2}{4}}}{2} \quad \textbf{(7.22.1)}$$

$V\_24 := subs(H=h, Ellipsoid\_V) + subs(H=h1, d=d1, Cone\_V)$

$$V\_24 := \frac{1}{6} \pi b c h + \frac{1}{12} \pi d1^2 h1 \quad \textbf{(7.22.2)}$$

#### 25. Cylinder + 3 Cones

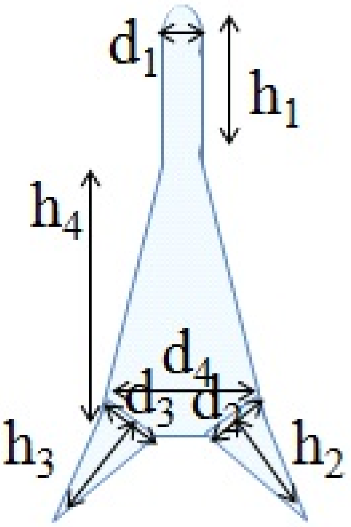


Area= MainConeSide + MainConeBase + TopCylinderSide +TopCylinderTop +BottomCone2Side

-BottomCone2Base+BottomCone3Side-BottomCone3Base

$A\_25 := subs(d2 = d4, d1 = d1, h = h4, Cone\_Trunc\_A\_Side) + subs(d = d4, Circle\_A) + subs(H = h1, d = d1, Cylinder\_Side\_A) + subs(d = d1, Circle\_A) + subs(H = h2, d = d2, Cone\_ASide) - subs(d = d2, Circle\_A) + subs(H = h3, d = d3, Cone\_ASide) - subs(d = d3, Circle\_A);$

$$A\_25 := \pi \left(\frac{d1}{2} + \frac{d4}{2}\right) \sqrt{\left(\frac{d4}{2} - \frac{d1}{2}\right)^2 + h4^2} + \frac{\pi\, d4^2}{4} + \pi\, d1\, h1 + \frac{\pi\, d1^2}{4} + \frac{\pi\, d2 \sqrt{h2^2 + \frac{d2^2}{4}}}{2} - \frac{\pi\, d2^2}{4} + \frac{\pi\, d3 \sqrt{h3^2 + \frac{d3^2}{4}}}{2} - \frac{\pi\, d3^2}{4} \quad \textbf{(7.23.1)}$$

$V\_25 := subs(d2 = d4, d1 = d1, h = h4, Cone\_Trunc\_V) + subs(H = h1, d = d1, Cylinder\_V) + subs(H = h2, d = d2, Cone\_V) + subs(H = h3, d = d3, Cone\_V);$

$$V\_25 := \frac{\pi\, h4\, (d1^2 + d1\, d4 + d4^2)}{12} + \frac{\pi\, d1^2\, h1}{4} + \frac{\pi\, d2^2\, h2}{12} + \frac{\pi\, d3^2\, h3}{12} \quad \textbf{(7.23.2)}$$

#### 27. Half sphere

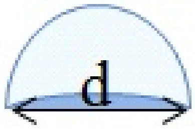


$V\_27 := \frac{Sphere\_V}{2}$

$$V\_27 := \frac{\pi\, d^3}{12} \quad \textbf{(7.24.1)}$$

$A\_27 := \frac{Sphere\_A}{2} + Circle\_A$

$$A\_27 := \frac{3\,\pi\,d^2}{4} \tag{7.24.2}$$

#### 34. 2 Half ellipsoids + Prism on elliptic base.

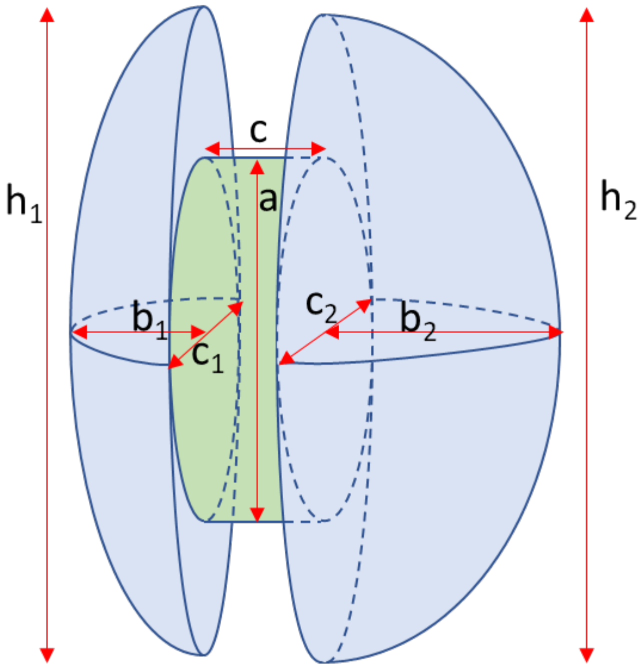


$$A\_34 := \frac{1}{2} subs(H=h1, b=2\cdot b1, c=c1, Ellipsoid\_A) + subs(a=h1, b=c1, Ellipse\_A) - subs(a=a, b=c1, Ellipse\_A) + \frac{1}{2} subs(H=h2, b=2\cdot b2, c=c2, Ellipsoid\_A) + subs(a=h2, b=c2, Ellipse\_A) - subs(a=a, b=c2, Ellipse\_A) + subs(a=a, b=c1, Ellipse\_P)\cdot c$$

$$A\_34 := \frac{\pi\,(2\,b1+c1)\left(b1+\frac{c1}{2}+\frac{2\,h1^2 \arcsin\left(\frac{\sqrt{4\,h1^2-(2\,b1+c1)^2}}{2\,h1}\right)}{\sqrt{4\,h1^2-(2\,b1+c1)^2}}\right)}{8} + \frac{\pi\,h1\,c1}{4} - \frac{\pi\,a\,c1}{4} + \frac{\pi\,(2\,b2+c2)\left(b2+\frac{c2}{2}+\frac{2\,h2^2 \arcsin\left(\frac{\sqrt{4\,h2^2-(2\,b2+c2)^2}}{2\,h2}\right)}{\sqrt{4\,h2^2-(2\,b2+c2)^2}}\right)}{8} + \frac{\pi\,h2\,c2}{4} - \frac{\pi\,a\,c2}{4} + \pi\left(\frac{a}{2}+\frac{c1}{2}\right)\left(1+\frac{(a-c1)^2}{4\,(a+c1)^2}\right)c \tag{7.25.1}$$

$$V\_34 := \frac{1}{2} subs(H=h1, b=2\cdot b1, c=c1, Ellipsoid\_V) + \frac{1}{2} subs(H=h2, b=2\cdot b2, c=c2, Ellipsoid\_V) + subs(a=a, b=c1, c=c, EllipticPrism\_V)$$

$$V\_34 := \frac{1}{6}\pi\, c1\, b1\, h1 + \frac{1}{6}\pi\, b2\, c2\, h2 + \frac{1}{4}\pi\, a\, c1\, c \tag{7.25.2}$$

### 35. Cymbelloid

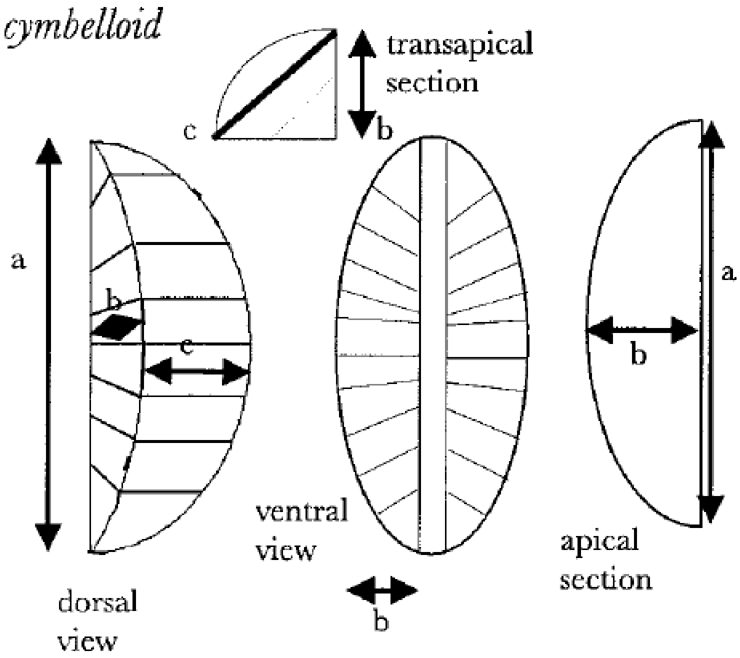


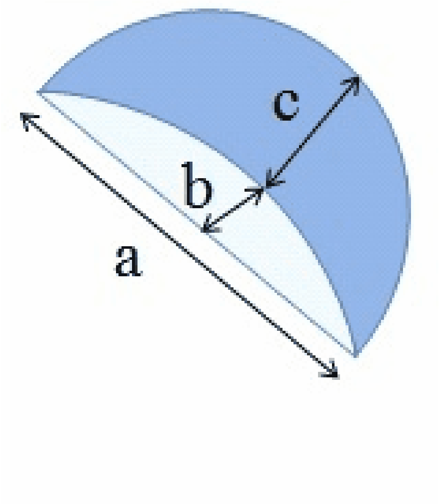


From Hi99 we find. ,

$\text{beta} := 2 \cdot \arcsin\left(\frac{c}{2 \cdot b}\right);$

$$\beta := 2 \arcsin\left(\frac{c}{2\, b}\right) \tag{7.26.1}$$

$$V\_35 := \frac{subs(H = a, b = 2 \cdot b, c = 2 \cdot b, Ellipsoid\_V) \cdot \text{beta}}{2 \cdot \text{pi}}$$

$$V\_35 := \frac{2\,a\,b^2 \arcsin\left(\frac{c}{2\,b}\right)}{3} \tag{7.26.2}$$

*subs*(*H* = *a*, *b* = 2 · *b*, *c* = 2 · *b*, *Ellipsoid_A*)

$$\pi\,b \left(2\,b + \frac{2\,a^2 \arcsin\left(\frac{\sqrt{4\,a^2 - 16\,b^2}}{2\,a}\right)}{\sqrt{4\,a^2 - 16\,b^2}}\right) \tag{7.26.3}$$

$A\_35 := subs(H = a, b = 2 \cdot b, c = 2 \cdot b, Ellipsoid\_A) \cdot \frac{\text{beta}}{2 \cdot \text{pi}} + 2 \cdot \frac{1}{2}\, subs(a = a, b = 2 \cdot b, Ellipse\_A)$

$$A\_35 := b \left(2\,b + \frac{2\,a^2 \arcsin\left(\frac{\sqrt{4\,a^2 - 16\,b^2}}{2\,a}\right)}{\sqrt{4\,a^2 - 16\,b^2}}\right) \arcsin\left(\frac{c}{2\,b}\right) + \frac{\pi\,a\,b}{2} \tag{7.26.4}$$

*Ellipsoid_KTA*

$$4\,\pi \left(\frac{\left(\frac{a}{2}\right)^p \left(\frac{b}{2}\right)^p}{3} + \frac{\left(\frac{a}{2}\right)^p \left(\frac{c}{2}\right)^p}{3} + \frac{\left(\frac{b}{2}\right)^p \left(\frac{c}{2}\right)^p}{3}\right)^{\frac{1}{p}} \tag{7.26.5}$$

*subs*(*H* = *a*, *b* = 2 · *b*, *c* = 2 · *b*, *Ellipsoid_KTA*)

$$4\,\pi \left(\frac{2\left(\frac{a}{2}\right)^p b^p}{3} + \frac{\left(b^p\right)^2}{3}\right)^{\frac{1}{p}} \tag{7.26.6}$$

$A\_35KT := subs(H = a, b = 2 \cdot b, c = 2 \cdot b, Ellipsoid\_KTA) \cdot \frac{\text{beta}}{2 \cdot \text{pi}} + 2 \cdot \frac{1}{2}\, subs(b = 2 \cdot b, Ellipse\_A)$

$$A\_35KT := 4 \left(\frac{2\left(\frac{a}{2}\right)^p b^p}{3} + \frac{\left(b^p\right)^2}{3}\right)^{\frac{1}{p}} \arcsin\left(\frac{c}{2\,b}\right) + \frac{\pi\,a\,b}{2} \tag{7.26.7}$$

#### 40. Gomphonemoid

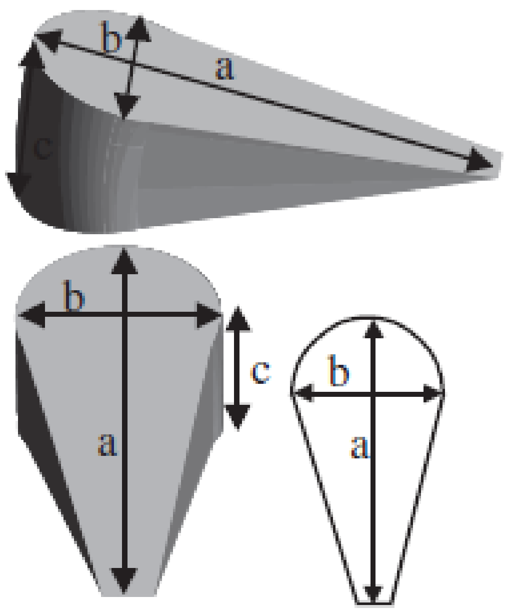


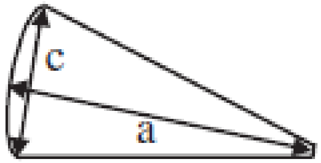


apical section girdle view

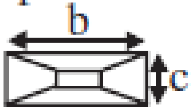


transapical view from base pole

apical section valve view

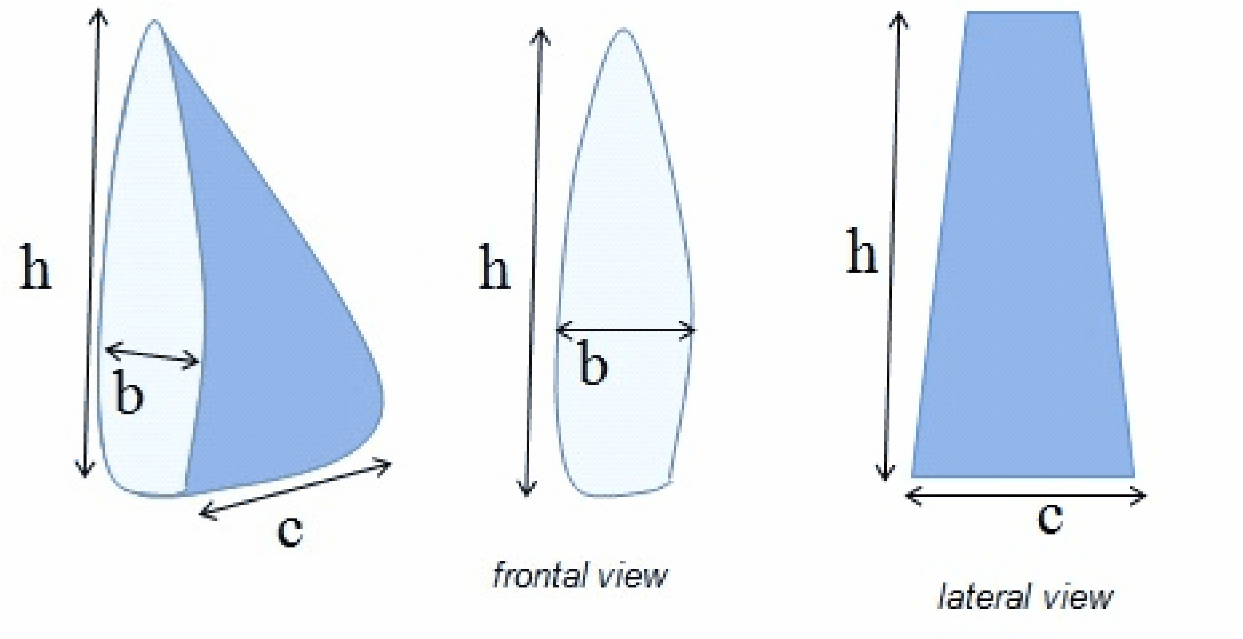


There is no definition of this shape. But such cells are rare. I use formulas from Sin 03, but the problem that in one case I get spherisity less than 1, which means that something is wrong with the formula. We use notation from ASh

$$\mathbf{A} \approx \frac{b}{2}\left[2h + \pi h \operatorname{asin}\left(\frac{c}{2h}\right) + \left(\frac{\pi}{2} - 2\right) b\right]$$

$A\_40 := \frac{b}{2}\left(2 \cdot h + \mathrm{pi} \cdot h \cdot \arcsin\left(\frac{c}{2\,h}\right) + \left(\frac{\mathrm{pi}}{2} - 2\right) \cdot b\right);$

$$A\_40 := \frac{b\left(2\,h + \pi\,h \arcsin\left(\frac{c}{2\,h}\right) + \left(\frac{\pi}{2} - 2\right) b\right)}{2} \qquad \textbf{(7.27.1)}$$

$$\mathbf{V} \approx \frac{\mathrm{hb}}{4}\left[\mathrm{h}+\left(\frac{\pi}{4}-1\right)\mathrm{b}\right]\operatorname{asin}\left(\frac{\mathrm{c}}{2\mathrm{h}}\right)$$

$V\_40 := \frac{h \cdot b}{4} \cdot \left(h + \left(\frac{pi}{4} - 1\right) \cdot b\right) \cdot \arcsin\left(\frac{c}{2 \cdot h}\right);$

$$V\_40 := \frac{h\, b \left(h + \left(\frac{\pi}{4} - 1\right) b\right) \arcsin\left(\frac{c}{2\, h}\right)}{4} \quad (7.27.2)$$

#### 41 Sickle-shaped prism

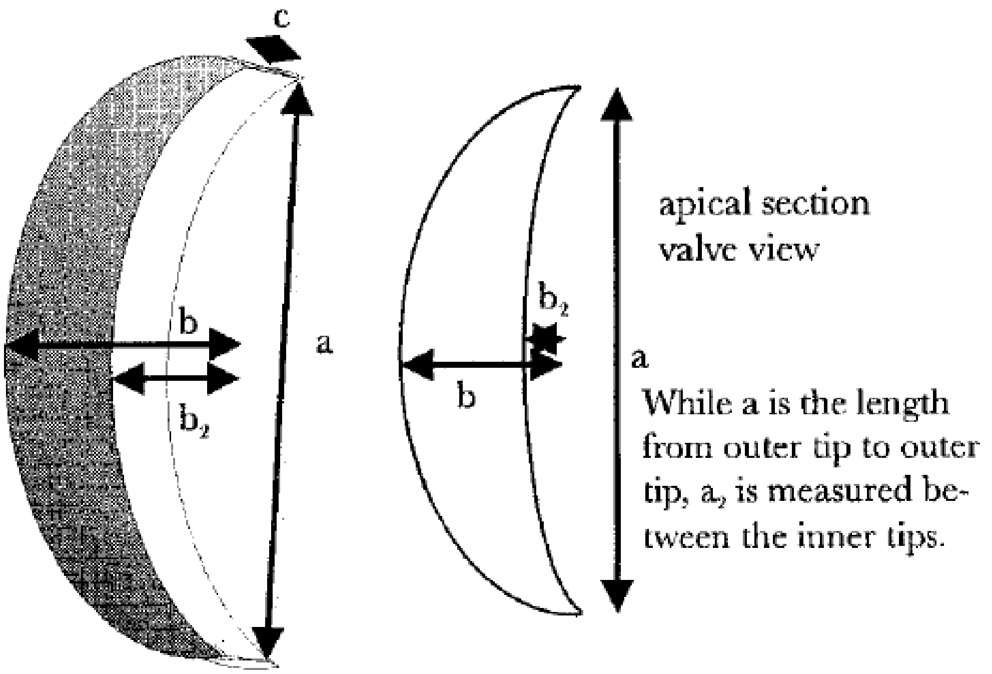


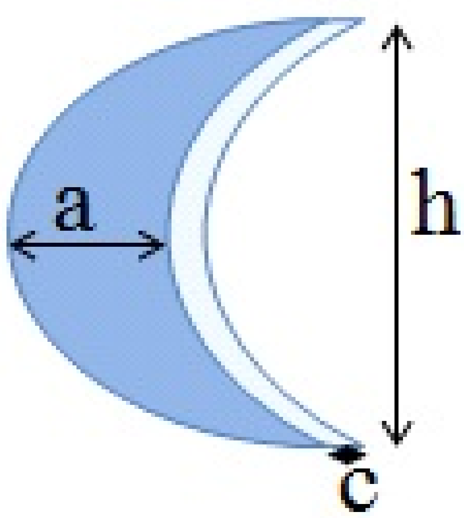


$V\_41 := factor\left(\frac{1}{2} subs(b = 2 \cdot b, a = h, EllipticPrism\_V) - \frac{1}{2} subs(b = 2 \cdot b2, a = h,\right.$

$$EllipticPrism\_V)\Big)$$

$$V\_41 := \frac{\pi\, c\, h\, (b - b2)}{4} \tag{7.28.1}$$

$$A\_41 := 2\cdot\left(\frac{1}{2}\cdot subs(b = 2\cdot b, a = h, Ellipse\_A) - \frac{1}{2}\cdot subs(b = 2\cdot b2, a = h, Ellipse\_A)\right) + \left(\frac{1}{2}\cdot subs(b = 2\cdot b, a = h, Ellipse\_P) + \frac{1}{2}\cdot subs(b = 2\cdot b2, a = h, Ellipse\_P)\cdot c\right)$$

$$A\_41 := \frac{\pi\, h\, b}{2} - \frac{\pi\, h\, b2}{2} + \frac{\pi\left(\frac{h}{2} + b\right)\left(1 + \frac{(h - 2\,b)^2}{4\,(h + 2\,b)^2}\right)}{2} + \frac{\pi\left(\frac{h}{2} + b2\right)\left(1 + \frac{(h - 2\,b2)^2}{4\,(h + 2\,b2)^2}\right)c}{2} \tag{7.28.2}$$

$$A\_41\_appr := \frac{\mathrm{pi}\, h\, b}{2} - \frac{\mathrm{pi}\, h\, b2}{2} + \frac{\mathrm{pi}\left(\frac{h}{2} + b\right)c}{2} + \frac{\mathrm{pi}\left(\frac{h}{2} + b2\right)c}{2}$$

$$A\_41\_appr := \frac{\pi\, h\, b}{2} - \frac{\pi\, h\, b2}{2} + \frac{\pi\left(\frac{h}{2} + b\right)c}{2} + \frac{\pi\left(\frac{h}{2} + b2\right)c}{2} \tag{7.28.3}$$

$$\frac{\mathrm{pi}}{2}\cdot\left(normal\left(\frac{A\_41\_appr}{\frac{\mathrm{pi}}{2}}\right)\right)$$

$$\frac{\pi\,(b\,c + h\,b + c\,b2 - h\,b2 + c\,h)}{2} \tag{7.28.4}$$

#### 43 Prism on elliptic base + 4 Cones

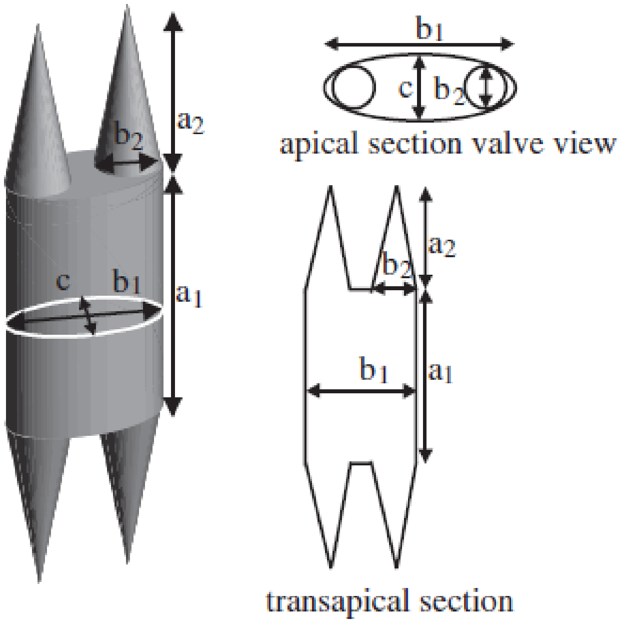


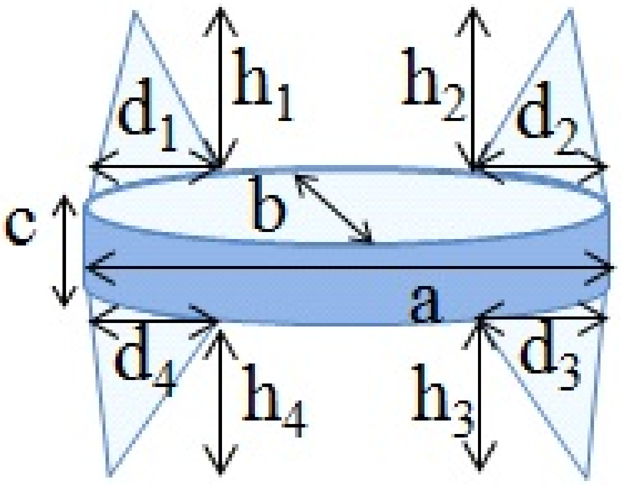


$A\_44 := EllipticPrism\_A - subs(d = d1, Circle\_A) - subs(d = d2, Circle\_A) - subs(d = d3, Circle\_A) - subs(d = d4, Circle\_A) + subs(H = h1, d = d1, Cone\_ASide) + subs(H = h2, d = d2, Cone\_ASide) + subs(H = h3, d = d3, Cone\_ASide) + subs(H = h4, d = d4, Cone\_ASide)$

$$A\_44 := c\,\pi\left(\frac{a}{2}+\frac{b}{2}\right)\left(1+\frac{(a-b)^2}{4\,(a+b)^2}\right)+\frac{\pi\,a\,b}{2}-\frac{\pi\,d1^2}{4}-\frac{\pi\,d2^2}{4}-\frac{\pi\,d3^2}{4}-\frac{\pi\,d4^2}{4}+\frac{\pi\,d1\sqrt{h1^2+\frac{d1^2}{4}}}{2}+\frac{\pi\,d2\sqrt{h2^2+\frac{d2^2}{4}}}{2}+\frac{\pi\,d3\sqrt{h3^2+\frac{d3^2}{4}}}{2}+\frac{\pi\,d4\sqrt{h4^2+\frac{d4^2}{4}}}{2} \quad (7.29.1)$$

Assuming that all h1..h4 are equal and d1..d4 are equal we obtain

$A\_44\_simpl := subs(h1 = h, h2 = h, h3 = h, h4 = h, d1 = d, d2 = d, d3 = d, d4 = d, A\_44);$

$$A\_44\_simpl := c\,\pi\left(\frac{a}{2}+\frac{b}{2}\right)\left(1+\frac{(a-b)^2}{4\,(a+b)^2}\right)+\frac{\pi\,a\,b}{2}-\pi\,d^2+2\,\pi\,d\sqrt{h^2+\frac{d^2}{4}} \quad \textbf{(7.29.2)}$$

$V\_44 := EllipticPrism\_V + subs(H=h1, d=d1, Cone\_V) + subs(H=h2, d=d2, Cone\_V) + subs(H=h3, d=d3, Cone\_V) + subs(H=h4, d=d4, Cone\_V)$

$$V\_44 := \frac{1}{4}\,\pi\,a\,b\,c+\frac{1}{12}\,\pi\,d1^2\,h1+\frac{1}{12}\,\pi\,d2^2\,h2+\frac{1}{12}\,\pi\,d3^2\,h3+\frac{1}{12}\,\pi\,d4^2\,h4 \quad \textbf{(7.29.3)}$$

### 44 Pyramid (rectangular base)

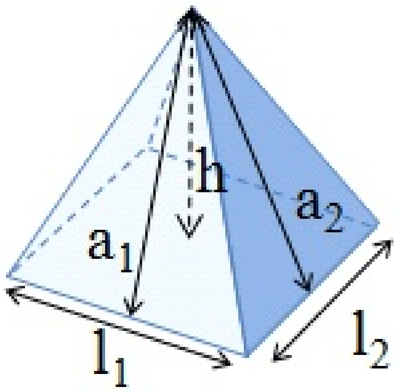


$l1 = l2 = d$

$$false \quad \textbf{(7.30.1)}$$

$A\_44 := d\cdot d + d\cdot \mathrm{sqrt}\left(h^2+\left(\frac{d}{2}\right)^2\right) + d\cdot \mathrm{sqrt}\left(h^2+\left(\frac{d}{2}\right)^2\right);$

$$A\_44 := d^2 + d\sqrt{d^2+4\,h^2} \quad \textbf{(7.30.2)}$$

$V\_44 := \frac{1}{3}\cdot d\cdot d\cdot h;$

$$V\_44 := \frac{d^2\,h}{3} \quad \textbf{(7.30.3)}$$

### 46. Prisma on triangle-base 2

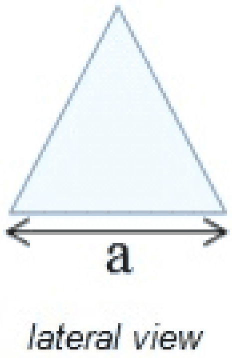


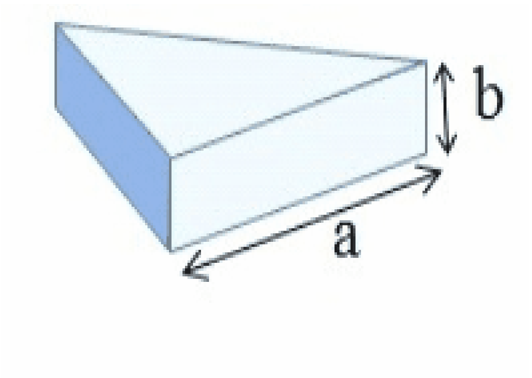


$$A\_46 := 3\cdot a\cdot b + 2\cdot 1/2\cdot a\cdot \frac{\text{sqrt}(3)}{2}\cdot a;$$

$$A\_46 := 3\,a\,b + \frac{\sqrt{3}\,a^2}{2} \qquad \textbf{(7.31.1)}$$

$$V\_46 := 1/2\cdot a\cdot \frac{\text{sqrt}(3)}{2}\cdot a\cdot b$$

$$V\_46 := \frac{a^2\sqrt{3}\,b}{4} \qquad \textbf{(7.31.2)}$$

#### 51. 2 Half ellipsoids

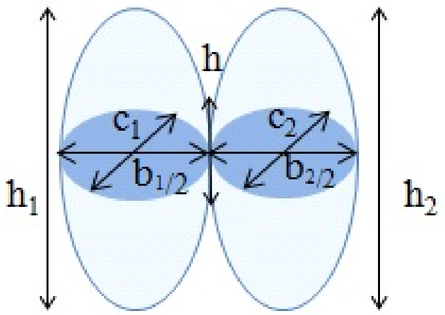


$h_1 = h_2$

$c_2 = c_1$

This formula assumes that h1=h2 and c1=c2, so two ellipses perfectly fit to each other

$$A\_51 := \frac{1}{2}\mathit{subs}(H=h1,\ b=2\cdot b1,\ c=c1,\ \mathit{Ellipsoid\_A}) + \frac{1}{2}\mathit{subs}(H=h2,\ b=2\cdot b2,\ c=c2,\ \mathit{Ellipsoid\_A})$$

$$A\_51 := \frac{\pi\,(2\,b1 + c1)\left(b1 + \frac{c1}{2} + \frac{2\,h1^2 \arcsin\left(\frac{\sqrt{4\,h1^2 - (2\,b1 + c1)^2}}{2\,h1}\right)}{\sqrt{4\,h1^2 - (2\,b1 + c1)^2}}\right)}{8} \qquad \textbf{(7.32.1)}$$

$$+\frac{\pi\,(2\,b2+c2)\left(b2+\frac{c2}{2}+\frac{2\,h2^{2}\arcsin\left(\frac{\sqrt{4\,h2^{2}-(2\,b2+c2)^{2}}}{2\,h2}\right)}{\sqrt{4\,h2^{2}-(2\,b2+c2)^{2}}}\right)}{8}$$

$V\_51 := \frac{1}{2} subs(H=h1, b=2\cdot b1, c=c1, Ellipsoid\_V) + \frac{1}{2} subs(H=h2, b=2\cdot b2, c=c2, Ellipsoid\_V)$

$$V\_51 := \frac{1}{6}\,\pi\,c1\,b1\,h1+\frac{1}{6}\,\pi\,b2\,c2\,h2 \qquad \textbf{(7.32.2)}$$
