## Supplementary material for "Shape matters: the relationship between cell geometry and diversity in phytoplankton": 38 geometric shapes

| Shape Class | Shape |
|---|---|
| Complex | 2 half ellipsoids + priism on elliptic base |
| Complex | Cylinder +2 cones |
| Complex | Cylinder+3cones |
| Complex | Ellipsoid+2cones+cylinder |
| Complex | Half ellipsoid + cone on elliptic base |
| Complex | Prism on elliptic base + 4 Cones |
| Complex | Prism on elliptic base+ box |
| Complex | prolate spheroid + 2 cylinder |
| Conic | Cone |
| Conic | cone+half sphere |
| Conic | Ellipsoid + cone |
| Conic | truncated cone + half sphere |
| Conic | 2 truncated cones |
| Conic | Double cone |
| Cylindrical | Cylinder |
| Cylindrical | Prism on elliptic base |
| Ellipsoidal (Spheric) | 2 half ellipsoids |
| Ellipsoidal (Spheric) | Cylinder +2 half spheres |
| Ellipsoidal (Spheric) | Ellipsoid |
| Ellipsoidal (Spheric) | Prolate spheroid |
| Ellipsoidal (Spheric) | rotational ellipsoid x 0.5 |
| Ellipsoidal (Spheric) | Sphere |
| Half shape | half cone |
| Half shape | half cone + cut flattened ellipsoid |
| Half shape | Half sphere |
| Other | 2 rotational ellipsoids |
| Other | 2 spheres * 5/8 |
| Other | Cymbelloid |
| Other | girdle diameter |
| Other | Gomphonemoid |
| Other | monoraphidioid |
| Other | Pyramid |
| Other | trapezoid |
| Prismatic | cube |
| Prismatic | Parallelepiped |
| Prismatic | parallelepiped/2 |
| Prismatic | Prism on parallelogram base |
| Prismatic | Prism on triangular base |
