## Supplementary material for "Shape matters: the relationship between cell geometry and diversity in phytoplankton": Fig. S

**Ryabov et al.**

### Supporting information

A

B

С

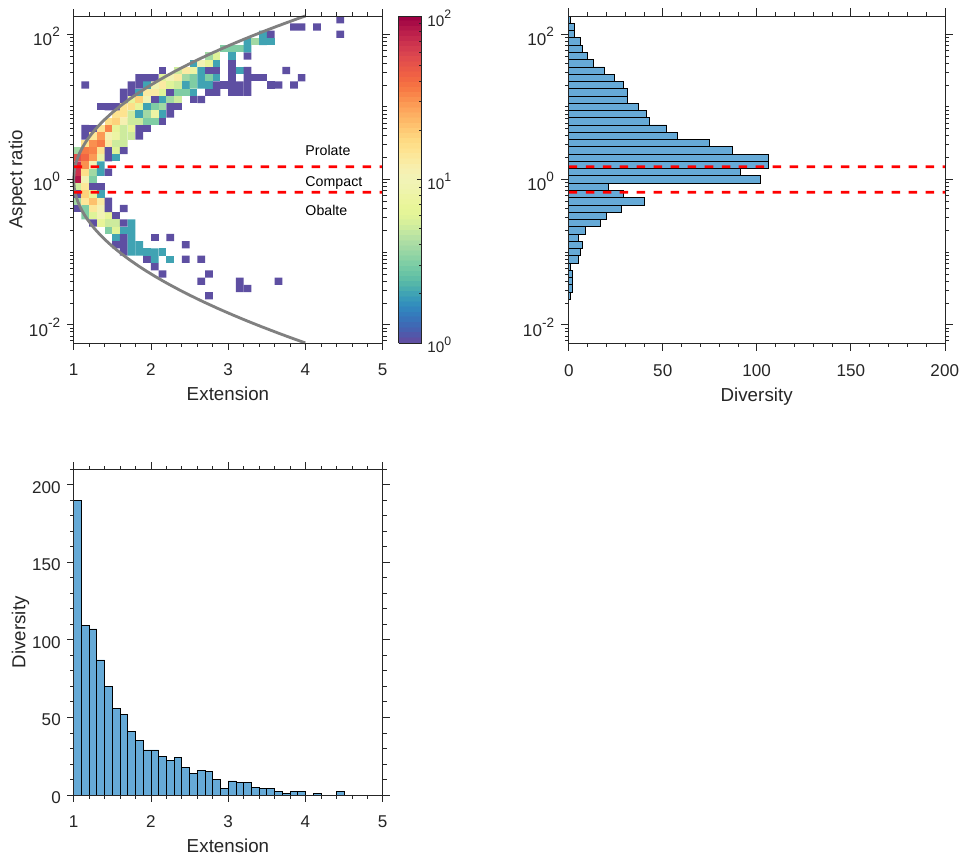

**Fig. S1. Bivariate effect of cell surface extension and aspect ratio on diversity.** (A) Distribution of taxonomic diversity (shown by colour) over aspect ratio (logarithmic binning) and surface extension. The grey line shows a fitting parabola $\log r=\pm1.3\sqrt{\varepsilon-1}$ to the upper boundary of the aspect ratio for a given surface extension. Horizontal red lines at $r=3/2$ and $r=2/3$ show the borders between compact, oblate and prolate cells, as defined in Methods. Diversity peaks for compact cells with smallest sphericity ($r=1, \varepsilon=1$) and decreases both with increasing surface extension and absolute value of logarithm of aspect ratio. (B) Distribution of taxonomic diversity over aspect ratio. When projected on this axis the distribution of diversity shows peaks for cells with $r=2$ and $r=1/2$. We suppose that these peaks occur due to the specific shape of the distribution in Fig. A, where aspect ratios of compact shapes can change very fast with a small increase in surface extension, so the distribution is strongly stretched in the vertical direction resulting in a local minimum at $r=1$. (C) Distribution of taxonomic diversity over surface extension. In this projection the diversity distribution decays exponentially.

A

B

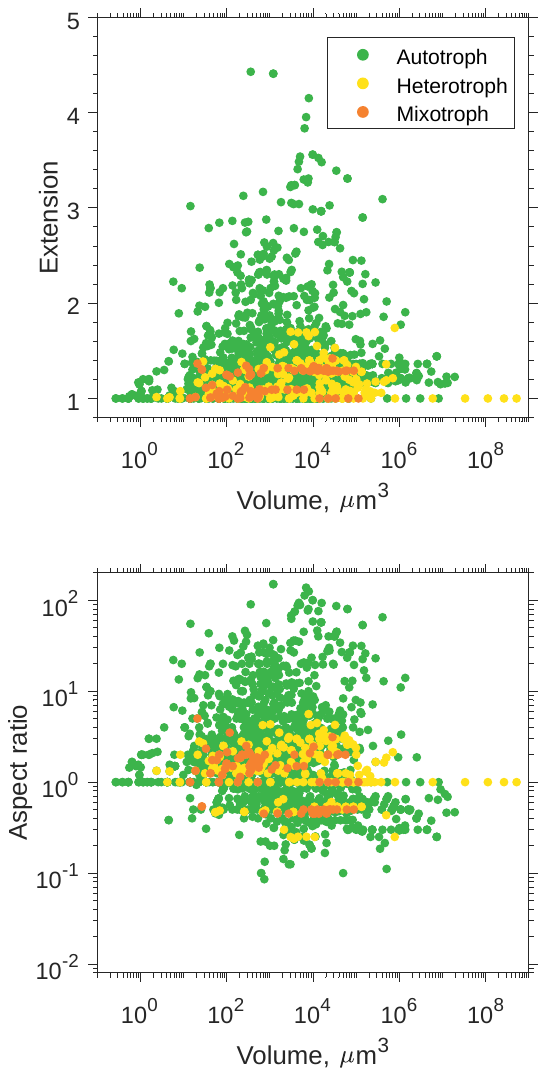

**Fig. S2. Dependence of the geometry of unicellular phytoplankton on cell volume for different nutritional modes.** (A) Surface extension and (B) aspect ratio for different heterotrophic groups of plankton cells (colour coded are autotrophs, mixotrophs and heterotrophs) in dependence of cell volume. Based on data from Baltic Sea, because only this data contained information on trophic levels of organisms.

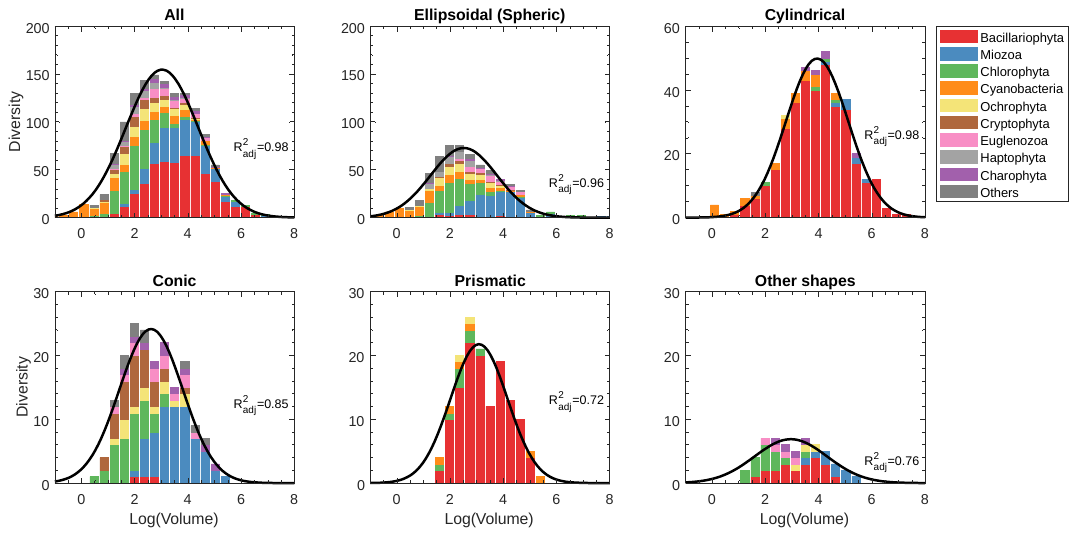

A

B

C

D

E

F

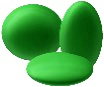

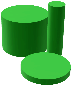

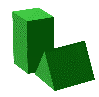

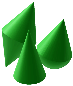

**Fig. S3. Distribution of taxonomic diversity as a function of volume for the most common shapes partitioned by phyla groups.** Black lines show a least square fit of a Gaussian function $D=a\exp\left( -\frac{\left( \log V-\log V_{0} \right)^{2}}{2\sigma^{2}} \right)$ to the histogram (see Table S1 for fitting parameters).

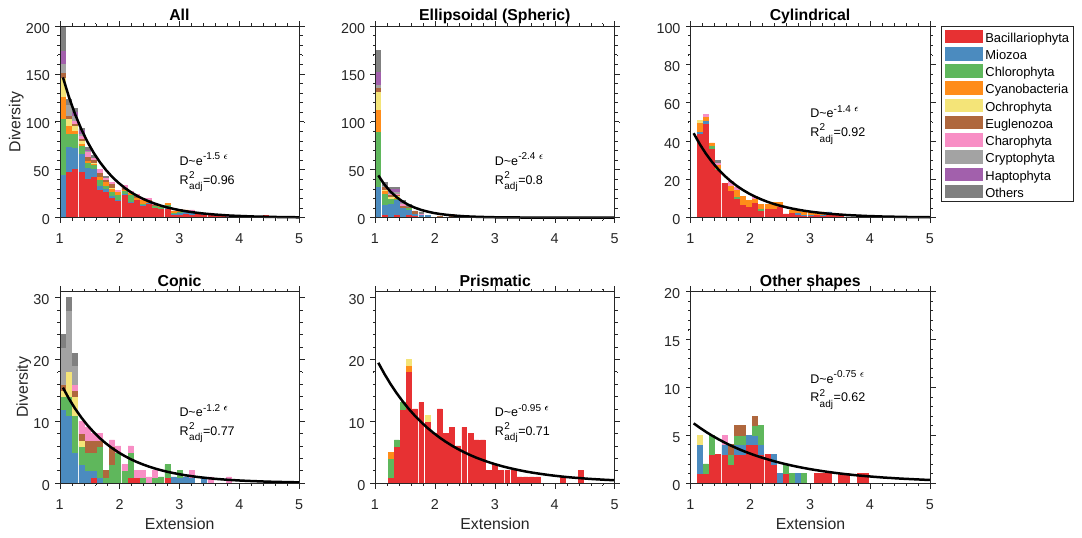

A

B

C

D

E

F

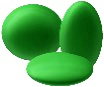

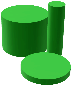

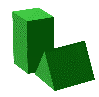

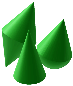

**Fig. S4. Distribution of taxonomic diversity as a function of surface extension for the most common shape types partitioned by phyla.** Solid lines show a least square fit of a linear function $\ln D=a-k\varepsilon$ to the log-transformed histogram (see Table S1 for fitting parameters).

How correlations obtained across all shapes ($R^{2}=0.96$) can be larger than those obtained for specific shapes? The diversity distribution of elliptical genera (Fig. B) abruptly decreases with $\varepsilon$ and elliptical genera have a strong tendency to be compact ($\varepsilon\approx1$). The maximum of diversity distribution for cylindrical and conic genera occurs at $\varepsilon\approx1.2$ (Fig. C,D). Finally, the diversity distributions of prismatic and other genera (Fig. E,F) decreases much slower with $\varepsilon$ and exhibit some secondary maxima. This can be interpreted as a kind of niche separation between shapes classes along the gradient of surface extension. This niche separation diminishes the quality of fit for each specific shape class, but it is not visible any more when we consider the entire distribution (Fig. A). The same explanation is applied for Fig. 6 (main text).

A

B

C

D

E

F

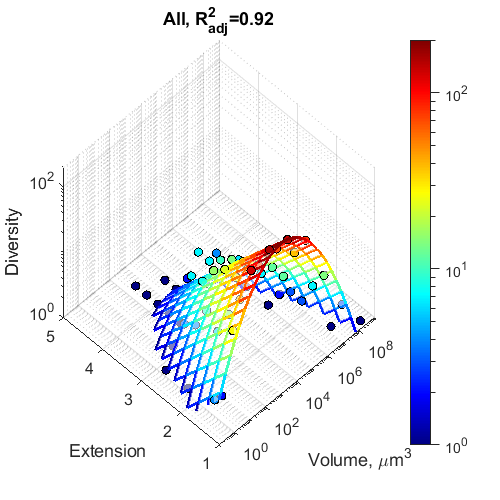

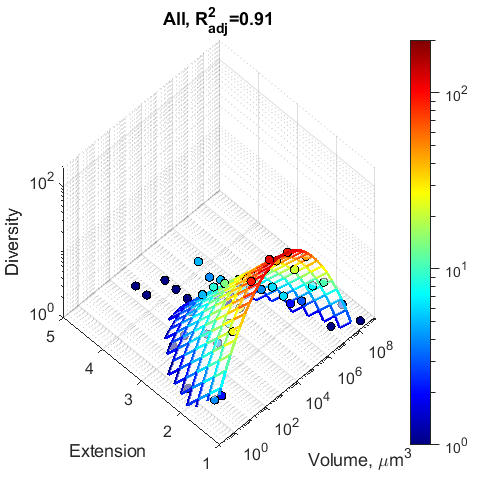

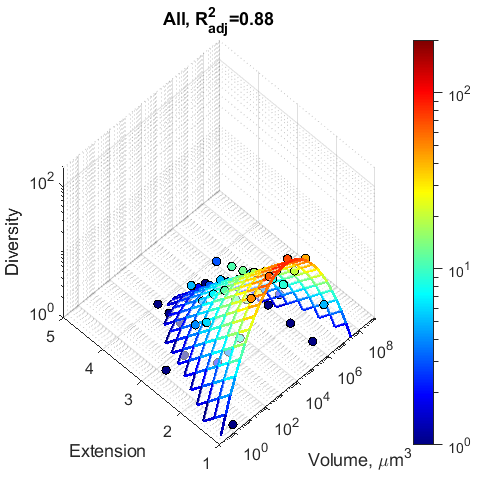

**Fig. S5. Comparison of biodiversity distribution calculated for all data (A), for the Baltic Sea data (B), and for the six ecoregions around the globe.** Shown are bivariate histograms of taxonomic diversity, D as a function of the surface extension and logarithm of cell volume. Fig. A is the same as Fig 6A in the main text and is shown here for better visual comparison.

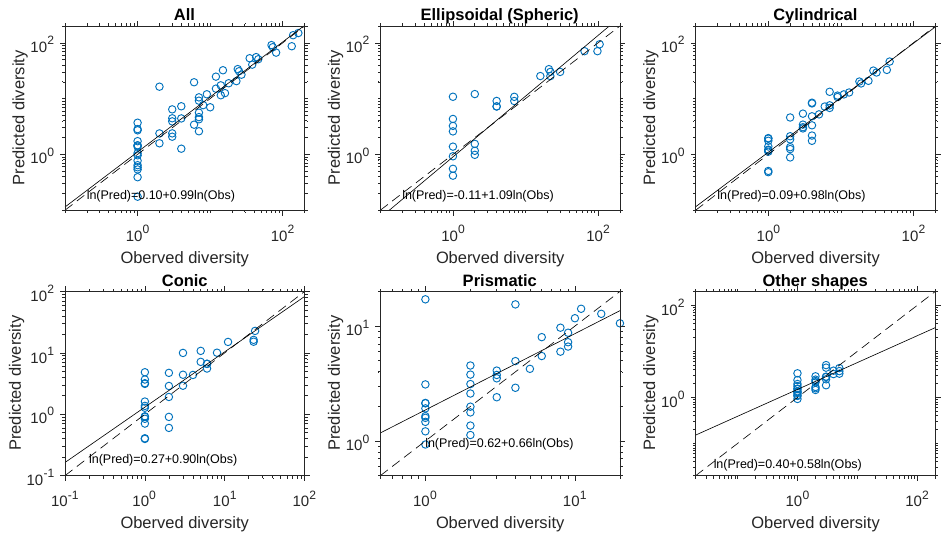

A

B

C

D

E

F

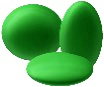

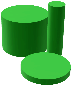

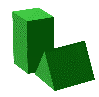

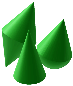

**Fig. S6. Comparison of observed diversity and diversity predicted based on nonlinear regression models in Fig. 6 (blue dots).** Black dashed lines show 1:1 diagonals and solid lines are linear regressions through the data points, see Table S1 for regression parameters. The closer the solid line is to the dashed line, and the smaller the variability of datapoints around this line, the better is the prediction of diversity by the model function $D=a\exp\left( -(log V-v_{0} \right)^{2}/(2 \sigma^{2})-k\varepsilon)$ in Fig. 6 (main text). An increase in the variation of the predicted diversity in the range of small $D$ can partly be explained by the fact that observed $D$ is constrained by 1, while predicted values can be less than 1. The regression analysis shows that the predictions for ellipsoidal (**B**), cylindrical (**C**) and conic (**D**) shapes are unbiased, because the solid and dashed lines are almost parallel. By contrast, for prismatic (**E**) and other shapes (**F**) the regression lines deviate from the diagonals, and the model is biased as regression line diverges from the main diagonal and predictions of diversity in the range of small observed $D$ are overestimated. However, as prismatic and other shapes are relatively rare, the model provides also a good and unbiased prediction of diversity across all shapes (**A**).

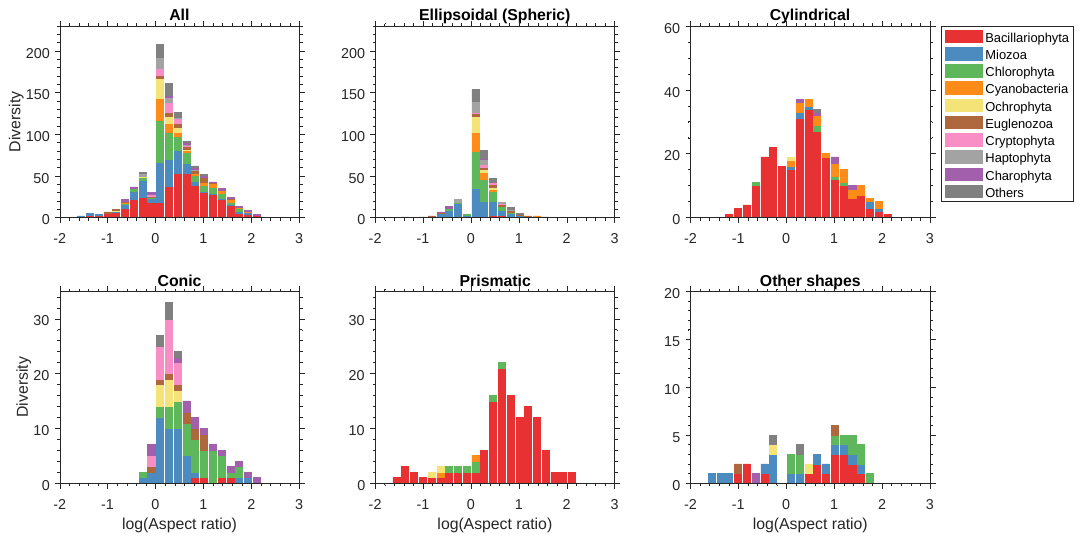

A

B

C

D

E

F

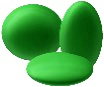

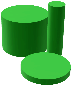

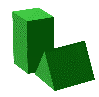

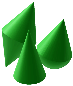

**Fig. S7. The same as in Fig. S4 but plotted as a function of the logarithm of the cell aspect ratio**.

| **Figure** | **Model** | $\boldsymbol{R}_{\boldsymbol{adj}}^{\boldsymbol{2}}$ | $\boldsymbol{b}_{\boldsymbol{1}}\boldsymbol{\pm}\boldsymbol{\delta}_{\boldsymbol{1}}\boldsymbol{(p)}$ | $\boldsymbol{b}_{\boldsymbol{2}}\boldsymbol{\pm}\boldsymbol{\delta}_{\boldsymbol{2}}\boldsymbol{(p)}$ | $\boldsymbol{b}_{\boldsymbol{3}}\boldsymbol{\pm}\boldsymbol{\delta}_{\boldsymbol{3}}\boldsymbol{(p)}$ | $\boldsymbol{b}_{\boldsymbol{4}}\boldsymbol{\pm}\boldsymbol{\delta}_{\boldsymbol{4}}\boldsymbol{(p)}$ | $\boldsymbol{b}_{\boldsymbol{5}}\boldsymbol{\pm}\boldsymbol{\delta}_{\boldsymbol{5}}\boldsymbol{(p)}$ |
| --- | --- | --- | --- | --- | --- | --- | --- |
| **Fig. S3A** | $D=b_{1}\exp\left( -\frac{\left( \log V-\log b_{2} \right)^{2}}{2\left( b_{3} \right)^{2}} \right)$ | 0.98 | 155±4 | 1100±100 | 1.36±0.04 |  |  |
| **Fig. S3B** |  | 0.96 | 73±3 | 330±40 | 1.25±0.06 |  |  |
| **Fig. S3C** |  | 0.98 | 50±1 | 8700±800 | 1.17±0.04 |  |  |
| **Fig. S3D** |  | 0.85 | 24±2 | 400±10 (${10}^{-3}$) | 1.19±0.1 |  |  |
| **Fig. S3E** |  | 0.72 | 22±2 (1.5e-05) | 1200±40 (${10}^{-2}$) | 1.07±0.2 (${10}^{-4}$) |  |  |
| **Fig. S3F** |  | 0.76 | 6.9±0.5 | 900±30 (${10}^{-2}$) | 1.36±0.2 (${10}^{-5}$) |  |  |
| **Fig. S4A** | $\ln D= b_{1} - b_{4}\varepsilon$ | 0.96 | 6.6±0.2 | 1.50±0.06 |  |  |  |
| **Fig. S4B** |  | 0.8 | 6.3±0.6 | 2.4±0.3 |  |  |  |
| **Fig. S4C** |  | 0.92 | 5.2±0.2 | 1.36±0.08 |  |  |  |
| **Fig. S4D** |  | 0.77 | 4.3±0.3 | 1.2±0.1 |  |  |  |
| **Fig. S4E** |  | 0.71 | 4.0±0.3 | 0.95±0.1 |  |  |  |
| **Fig. S4F** |  | 0.62 | 2.6±0.3 | 0.75±0.1 |  |  |  |
| **Fig. S6A** | $\ln D_{pred}= b_{1}+ b_{2}\ln D_{obs}$ | 0.85 | 0.1±0.1 (0.44) | 0.99±0.05 |  |  |  |
| **Fig. S6B** |  | 0.66 | -0.1±0.3 (0.74) | 1.2±0.2 |  |  |  |
| **Fig. S6C** |  | 0.87 | 0.1±0.1 (0.41) | 0.98±0.06 |  |  |  |
| **Fig. S6D** |  | 0.65 | 0.3±0.2 (0.11) | 0.9±0.1 |  |  |  |
| **Fig. S6E** |  | 0.54 | 0.6±0.1 (1e-04) | 0.6±0.1 |  |  |  |
| **Fig. S6F** |  | 0.54 | 0.4±0.08 (5e-05) | 0.6±0.1 |  |  |  |
| **Fig. 7A** | $\ln D=-\frac{\left( \log V-b_{1} \right)^{2}}{2 \left( b_{2} \right)^{2}}+b_{3}-b_{4}r$ | 0.89 | 3.09±0.06 | 1.37±0.04 | 5.2±0.07 | 1.58±0.09 |  |
| **Fig. 7B** |  | 0.86 | 2.7±0.1 | 1.47±0.07 | 4.8±0.1 | 3.18±0.3 |  |
| **Fig. 7C** |  | 0.75 | 3.76±0.08 | 1.26±0.06 | 4.2±0.1 | 1.39±0.1 |  |
| **Fig. 7D** |  | 0.76 | 2.5±0.1 | 1.4±0.1 | 3.5±0.1 | 1.78±0.2 |  |
| **Fig. 7E** | $\ln D=-\frac{\left( \log V-b_{1} \right)^{2}}{2 \left( b_{2} \right)^{2}}+b_{3}-\frac{\left( \log r-b_{4} \right)^{2}}{2 \left( b_{5} \right)^{2}}$ | 0.66 | 3.18±0.09 | 1.03±0.08 | 2.7±0.1 | 0.91±0.05 | 0.56±0.05 |
| **Fig. 7F** |  | 0.23 | 2.3±0.7 (0.0037) | 2.1±0.7 (${10}^{-2}$) | 1.7±0.2 | 1.0±0.1 | 0.6±0.2 (${10}^{-4}$) |

**Table S1. Fitting parameters for Fig. 7 (main text) and Supplementary figures.** Parameter values are specified with standard error, p-value in brackets is shown only when $p>{10}^{-5}$.
